# Library docking for Cannabinoid-2 Receptor ligands

**DOI:** 10.64898/2026.03.19.713017

**Authors:** Moira M. Rachman, Christos Iliopoulos-Tsoutsouvas, Michael D. Sacco, Xinyu Xu, Cheng-Guo Wu, Emma Santos, Isabella S. Glenn, Lu Paris, Michelle K. Cahill, Suthakar Ganapathy, Tia A. Tummino, Yurii S. Moroz, Dmytro S. Radchenko, Meri Okorie, Vivianne Tawfik, John J. Irwin, Alexandros Makriyannis, Georgios Skiniotis, Brian K. Shoichet

**Affiliations:** Department of Pharmaceutical Chemistry, University of California San Francisco, 1700 4th St., Byers Hall Rm 508D, San Francisco, CA 94143; Center for Drug Discovery and Department of Pharmaceutical Sciences, Northeastern University, Boston, MA 02115, USA; Department of Molecular and Cellular Physiology, Stanford University School of Medicine, Stanford, CA 94305, USA; Department of Anesthesiology, Perioperative and Pain Medicine, Stanford University School of Medicine, Stanford CA 94305 USA; Enamine Ltd., 67 Winston Churchill Street, Kyiv 02094, Ukraine; National Taras Shevchenko University of Kyiv, 60 Volodymyrska Street, Kyiv 01601, Ukraine; Chemspace LLC, 85 Winston Churchill Street, Suite 1, Kyiv 02094, Ukraine; Graduate Program in Pharmaceutical Sciences and Pharmacogenomics, University of California, San Francisco, San Francisco, CA 94158, USA; Department of Chemistry and Chemical Biology, Northeastern University, Boston, MA 02115, USA; Department of Structural Biology, Stanford University School of Medicine, Stanford, CA 94305, USA; Department of Structural Biology & Center of Excellence for Structural Cell Biology, St. Jude Children’s Research Hospital, Memphis, TN 38105, USA

## Abstract

Cannabinoid receptors are therapeutically promising GPCRs that are also interesting test systems for structure-based methods, which have targeted them previously. Here we used the CB2 receptor as a template to explore several topical questions in library docking. Whereas an earlier campaign against the CB1 receptor led to potent but relatively non-selective ligands, here we found that targeting interactions with polar, orthosteric site residues led to subtype-selective ligands. Docking hit rate and especially hit affinity improved in moving from a 7 million to a 2.6 billion molecule library. Similar to earlier studies, docking against active and inactive states of the receptor did not reliably bias toward the discovery of agonists or inverse agonists. Cryo-EM structures of two of the new agonists, each in a different chemotype, superposed well on the docking predictions. Correspondingly, structure-based optimization led to 10- to 140-fold improvements within three different series, also consistent with well-behaved ligand families. Hit rates with a fully enumerated 2.6 billion molecule library resembled those of an implied 11 billion molecule library from a building-block method, consistent with the latter’s ability to explore this space, though higher affinities were discovered from the fully enumerated set. Overall, eight diverse families of ligands, with potencies <100 nM and mostly unrelated to previously known ligands were found. Implications for future studies are considered.

## INTRODUCTION

The two primary cannabinoid receptors, CB1 and CB2, are lipid-family G protein-coupled receptors (GPCRs) that have attracted attention for their role in nociception, consciousness, homeostatic temperature regulation, satiety, and inflammatory responses^1–5^. The CB1 subtype is principally responsible for the CNS-related functions of endogenous agonists like anandamide and 2-arachidonoylglycerol, while the CB2 subtype is typically expressed peripherally where it modulates immune and inflammatory response. These roles have inspired decades of effort to exploit the receptors as therapeutic targets in analgesia, metabolic diseases, inflammation and immune response^6^, supporting the development of well-behaved functional assays^7^, the determination of atomic resolution structures in activated and inactive states^8–12^, and the development of tool molecules and clinical candidates^13–18^, including by library docking^19, 20^.

The many tools available to probe the cannabinoid receptors, and the many structures determined for them, has made them akin to model systems for structure-based discovery^21^. These have explored the impact of new methods to represent large chemical libraries and how the physical chemistry of those libraries affects docking success^22^, while other efforts have sought to discover new chemical tools and therapeutic leads for the CB1 and CB2 receptors^23, 24^, including mechanistic studies^25–29^.

Drawing on this body of work, we explore several topical questions in large library docking, using the CB2 receptor as a template. We investigated: **i.** Strategies to find ligands selective for the CB2 versus the CB1 receptor, to which it has 44% sequence identity (∼68% in the orthosteric site, and up to ∼90% considering only binding site residues). This was allied with efforts to retain favorable ligand physical properties, a challenge in lipid receptors. **ii.** We investigated whether we could bias ligand discovery toward agonists and inverse agonists by targeting the active and inactive states of the receptor, as captured by cryo-EM studies^8, 30^. This has typically been problematic for docking, with studies often finding a mix of ligand functions irrespective of the receptor conformation. Here we wondered if functionally-selective ligands would be more readily found in ultra-large libraries, which might be more likely to have molecules more specific for each receptor state than smaller libraries. **iii.** Leveraging these studies, we wanted to compare how docking hit rates and hit affinities would change with larger versus smaller molecules. Earlier studies have suggested that, with the better and deeper exploration of chemotypes in larger libraries, hit rates and affinities improve with library size^31^. How true this would be in the challenging site of a lipid receptor was unclear. **iv**. We also wanted to compare docking a fully enumerated large library—here 2.6 billion molecules, explicitly docked—with the results of docking an 11.5 billion implied molecules against the same form of the CB2 receptor^19^. **v.** Finally, with hundreds of molecules tested, we were interested in the diversity of molecules appearing from this structure-based study to the chemotypes extant in the ChEMBL^32^ and IUPHAR libraries. To support these studies, concentration-response curves were measured and key chemotypes were advanced in structure-activity relationship (SAR) series to 10- to 140-fold improved potency, and two cryo-EM structures were determined.

## RESULTS

### Selectivity-driving interactions in docking to the CB2 receptor

We began with a CB2 receptor structure (6PT0^30^) in its activated state. The structure’s ability to prioritize CB2R agonists through docking^33^ was tested with known ligands in the IUPHAR database^34^. Physically similar but topologically distinct molecules were used as decoys, using the DUDE-Z tools^35^. Although, the model differentiated actives from decoys well (logAUC: 17.5, **SI Figure S1**), the docking did not distinguish between selective and non-selective ligands, something also observed in earlier docking to CB1R^22^, which shares up to ∼90% sequence identity of binding site residues (**Figure 1A**), and even the non-conserved residues resemble each other physically (**Figure 1B**). Still, we noted that low lipophilicity correlates with CB2R selectivity (**Figure 1C**)^26^ and so sought to reduce the hydrophobicity of high-ranking docked molecules by prioritizing polar interactions in the otherwise lipophilic site. To do so, we increased the local dipoles of polar residues including T3.33, S7.39, S2.60, H.265 and the backbone of L182^ECL2^ (Ballesteros numbering), without changing their net charge, in our electrostatics calculations, as in earlier studies^36–38^. This raised the magnitude of the electrostatic component of the DOCK3.8 score (**SI Figure S2**). Post-docking filters for modeled hydrogen-bonds with these residues were also used.

**Figure 1.**
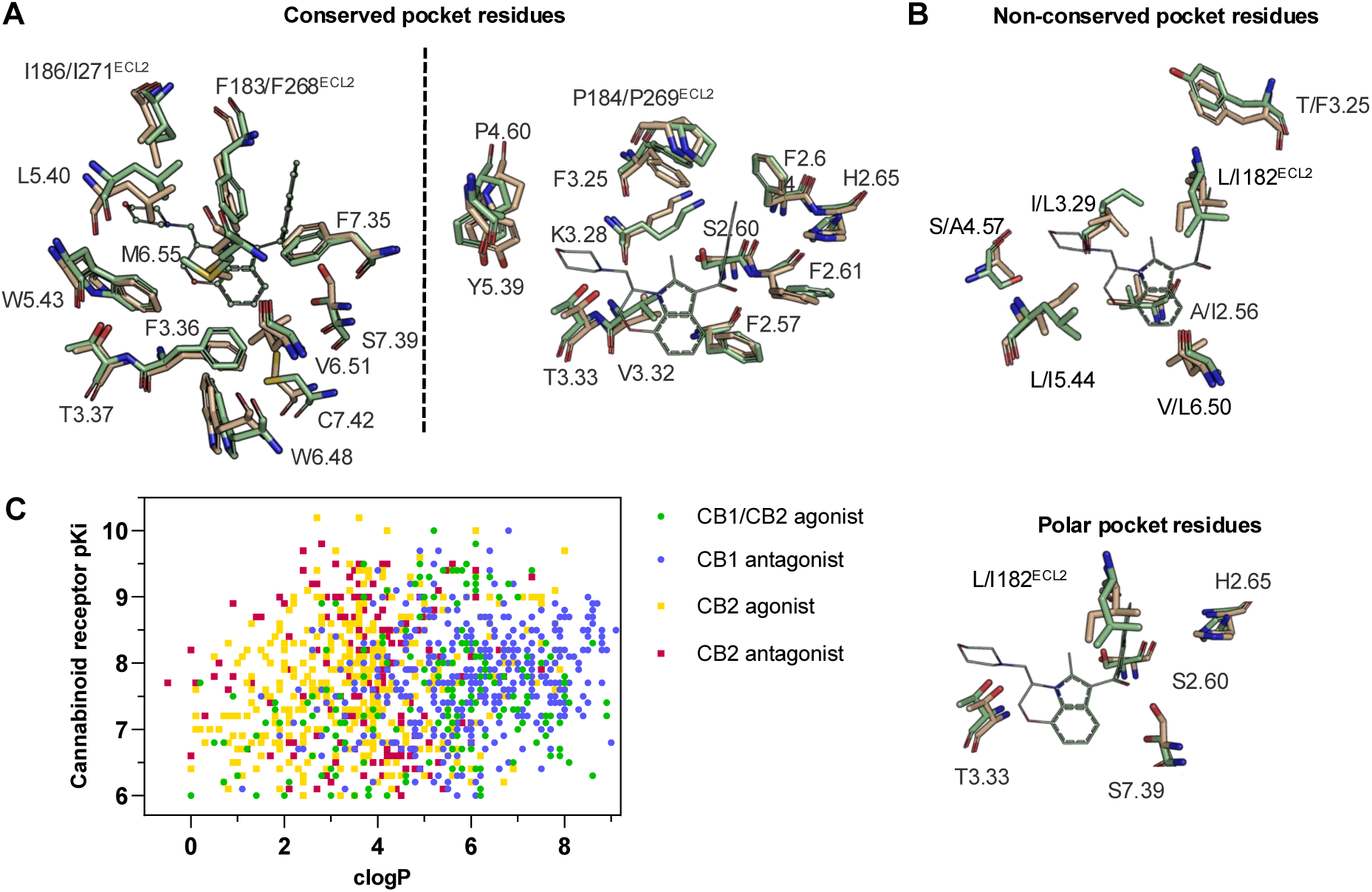
CB2R(green)/CB1R(tan) conserved pocket residues (**A**) and pocket residues the two subtypes do not have in common (**B**). Residues are displayed using Ballesteros-Weinstein numbering. Affinity of literature cannabinoid receptor ligands against their calculated lipophilicity (clogP), and polar pocket residues that were targeted for CB2R selectivity (**C**). Compounds reported with nanomolar affinities published between 1993 and 2023 in J. Med. Chem., Bioorg. Med. Chem., Bioorg. Med. Chem. Lett., ACS Med. Chem. Lett., ChemMedChem, RSC Med. Chem., and Eur. J. Med. Chem. were included. A compound was defined as selective if a 100-fold difference was reported. clogP was calculated using ChemAxon’s Chemicalize.

With the energy grids used for scoring thus parameterized, we docked a relatively small in-stock library of about 7 million molecules against the CB2 receptor. Each library molecule was sampled in and average of 1,720 orientations and 335 conformations, or about 576,200 configurations each in the receptor’s orthosteric site. After filtering for conformational strain^39^, 1.4M molecules with DOCK3.8 scores better than −20 remained. Filtering for polar interactions and excluding molecules with a hydrogen-bond donor that lacked a complementary acceptor in the orthosteric site, or that had more than 3 hydrogen-bond acceptors without a complementary receptor donor, left 101,559 molecules. These were filtered for novelty to 341 known cannabinoid receptors ligands from the ChEMBL database^32^, stipulating Tanimoto coefficients (Tc) ≤0.35 using ECFP4 fingerprints (a measurement of topological similarity, with values ≤0.35 typically indicating a scaffold hop for this fingerprint^40, 41^). The 100,543 molecules that remained were clustered for diversity (Tc ≤0.35), and after visual inspection 34 of these in-stock molecules were purchased for testing in vitro (**Methods**).

These 34 molecules were tested for binding by radioligand ([^3^H]-CP55,940) displacement at 10 μM, with eight displacing ≥30% (**Supporting Information In-Stock_Screen.xlsx**). These were checked for colloidal aggregation: two formed particles by dynamic light scattering and inhibited the counter-screening enzyme MDH in a detergent-dependent manner^42^, suggesting that they were acting as aggregators; these were no longer pursued (**SI Figure S3**). The six remaining molecules were tested in full concentration-response, revealing apparent K_i_ values ranging from 0.9 to 7.1 µM by Cheng-Prusoff^43^ (**Table 1**, **SI Figure S4**), an overall hit-rate of 17% (hit-rate = number active/number tested). Four of these were further tested for activity on CB1, and while ‘8876 and ‘1138 showed little selectivity between the sub-types, ‘6138 and ‘7302 were at least ∼40-fold selective for the CB2 over the CB1 receptor (**Table 1**, **SI Figure S4**). This was in contrast to our earlier docking against the CB1 receptor, which ultimately led to potent ligands, few of which were subtype selective.

**Table 1.**
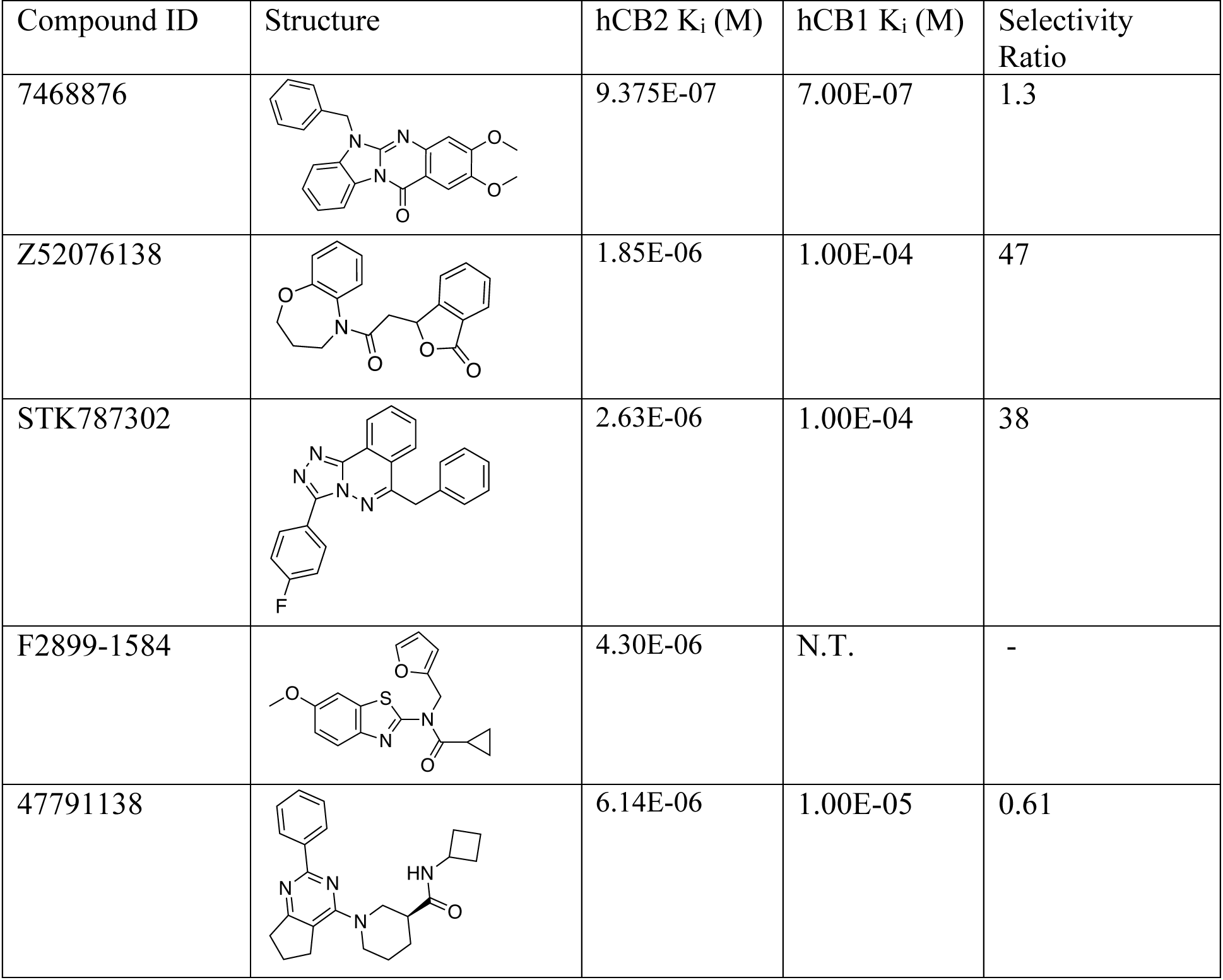
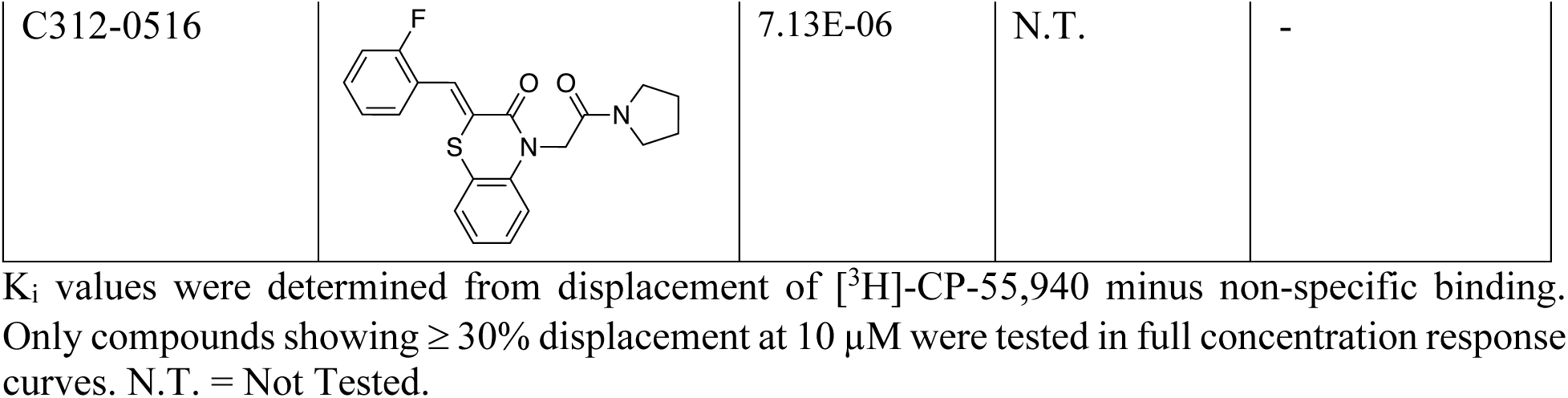
K_i_ values and CB1 selectivities from the 7 million library docking.

| Compound ID | Structure | hCB2 $K_i$ (M) | hCB1 $K_i$ (M) | Selectivity Ratio |
| --- | --- | --- | --- | --- |
| 7468876 |  | 9.375E-07 | 7.00E-07 | 1.3 |
| Z52076138 |  | 1.85E-06 | 1.00E-04 | 47 |
| STK787302 |  | 2.63E-06 | 1.00E-04 | 38 |
| F2899-1584 |  | 4.30E-06 | N.T. | - |
| 47791138 |  | 6.14E-06 | 1.00E-05 | 0.61 |
| C312-0516 |  | 7.13E-06 | N.T. | - |

$K_i$ values were determined from displacement of [ $^3\text{H}$ ]-CP-55,940 minus non-specific binding. Only compounds showing $\geq 30\%$ displacement at $10\ \mu\text{M}$ were tested in full concentration response curves. N.T. = Not Tested.

Encouraged by this in-stock screen, we moved to docking a larger 1.6 billion molecule library against the same activated structure of CB2. Because of the much larger library, we reduced sampling and increased the stringency of the score cutoff to −25. Only 143,500 configurations of each library molecule were sampled on average, rather than the 576,200 sampled in the in-stock screen. This allowed this screen to complete in 128,444 CPU hours, or 86 wall-clock hours on 1500 CPUs. After filtering out strained molecules about 22 million molecules remained. These were further filtered by the same interactions as previously, and clustered for diversity (see **Methods**). Visual examination of top-scoring cluster heads prioritized 197 molecules for synthesis and testing of which 151 were successfully made and purified by make-on-demand synthesis at Enamine from the REAL database (https://enamine.net/compound-collections/real-compounds) and at WuXi (https://chemistry.wuxiapptec.com/small_molecule), a fulfillment rate of 79%.

The 151 molecules were screened by radio-ligand displacement at single point, and 49 had ≥ 30% displacement (**Supporting Information 1.6B_Screen.xlsx**). Of these, 22 were tested in full concentration response and 16 had K_i_ values ≤ 15 μM by Cheng-Prusoff^43^ (**Table 2, SI Figure S5, S6A, S6B**). Most of these molecules, too, were selective against the CB1 receptor, with three having subtype selectivity of at least two-fold and nine more having selectivity of five-fold or better (**SI Figure S5, S6C**). These results, too, were consistent with the focus on polar interactions in the CB2R site leading to initial sub-type selectivity.

**Table 2.**
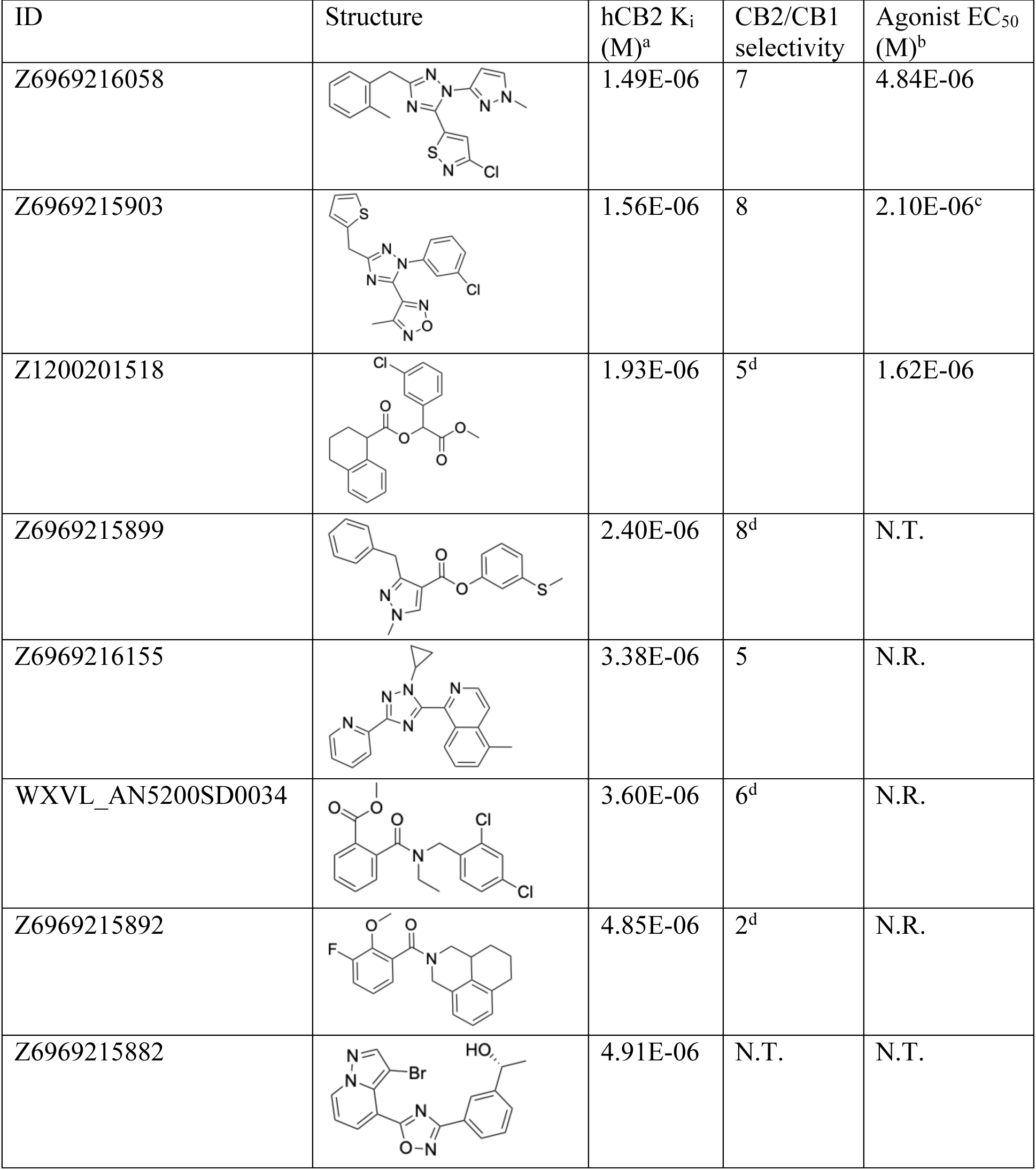
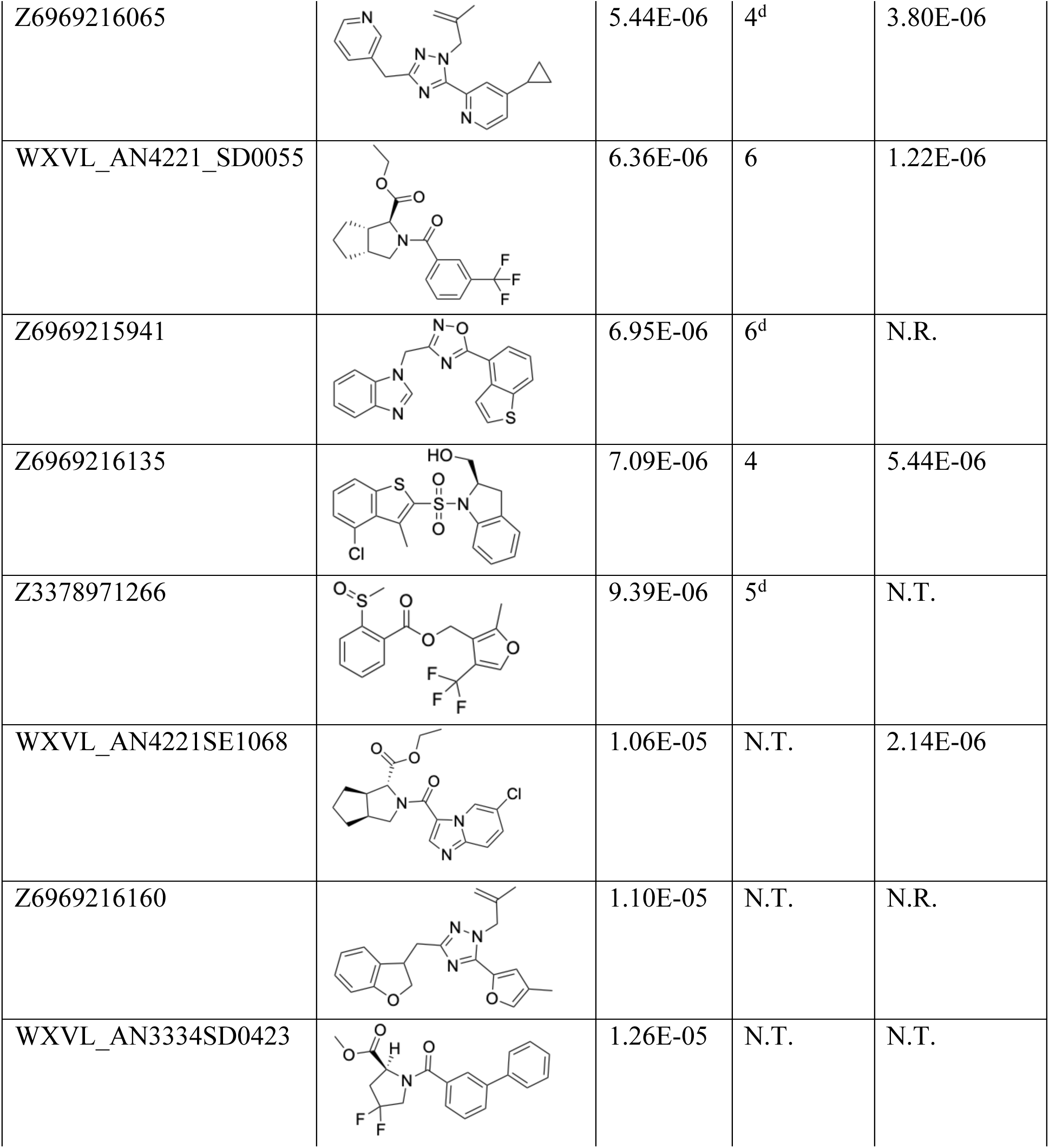
CB2R active ligands from a 1.6 billion library docking campaign. **a.** Ki values were determined from displacement of [^3^H]-CP-55,940 minus non-specific binding. Only compounds ≥ 30% displacement were tested in full concentration response curves. **b.** Agonism was determined by monitoring cAMP accumulation with GloSensor in the presence of Forskolin normalized to CP-55,940. **c.** Inverse agonism was determined by monitoring cAMP accumulation with GloSensor in the presence of Forskolin and CP-55,940 normalized to SR114528. **d.** CB1 K_i_‘s estimated by testing at 10 μM, 30 μM and 100 μM. N.R. = No Response. N.T. = Not Tested.

| ID | Structure | hCB2 K<sub>i</sub> (M)<sup>a</sup> | CB2/CB1 selectivity | Agonist EC<sub>50</sub> (M)<sup>b</sup> |
| --- | --- | --- | --- | --- |
| Z6969216058 |  | 1.49E-06 | 7 | 4.84E-06 |
| Z6969215903 |  | 1.56E-06 | 8 | 2.10E-06<sup>c</sup> |
| Z1200201518 |  | 1.93E-06 | 5<sup>d</sup> | 1.62E-06 |
| Z6969215899 |  | 2.40E-06 | 8<sup>d</sup> | N.T. |
| Z6969216155 |  | 3.38E-06 | 5 | N.R. |
| WXVL_AN5200SD0034 |  | 3.60E-06 | 6<sup>d</sup> | N.R. |
| Z6969215892 |  | 4.85E-06 | 2<sup>d</sup> | N.R. |
| Z6969215882 |  | 4.91E-06 | N.T. | N.T. |
| Z6969216065 |  | 5.44E-06 | 4<sup>d</sup> | 3.80E-06 |
| WXVL_AN4221_SD0055 |  | 6.36E-06 | 6 | 1.22E-06 |
| Z6969215941 |  | 6.95E-06 | 6<sup>d</sup> | N.R. |
| Z6969216135 |  | 7.09E-06 | 4 | 5.44E-06 |
| Z3378971266 |  | 9.39E-06 | 5<sup>d</sup> | N.T. |
| WXVL_AN4221SE1068 |  | 1.06E-05 | N.T. | 2.14E-06 |
| Z6969216160 |  | 1.10E-05 | N.T. | N.R. |
| WXVL_AN3334SD0423 |  | 1.26E-05 | N.T. | N.T. |

### Functional selectivity from the docking

In GPCR ligand discovery, one is typically interested in not only binders but in agonists, inverse agonists or antagonists. Docking’s success with such functional selectivity has been hit-and-miss, with some studies suggesting that docking against activated and inactivated states can bias toward agonists and inverse agonists, respectively^44, 45^ and many others finding a mix of agonists and inverse agonists irrespective of receptor conformation^46–49^. We wondered if moving to larger libraries, potentially including agonists and inverse agonists that better complemented the receptor, would improve functional selectivity for the CB2 receptor. Consistent with this hypothesis, whereas no agonists were found among the docking hits from 7 million molecule library, of the 12 large library hits that were tested for function, six were CB2R agonists with low µM EC_50_‘s (**Table 2**, **SI Figure S7**), and a seventh was an inverse agonist (**Table 2**, **SI Figure S8**). A caveat is that relatively few molecules were tested and the sampling in the large library campaign was relatively low, potentially missing important chemotypes.

If docking against an activated receptor conformation led to mostly agonists, docking against an inactive receptor conformation should lead to inverse agonists. To test this idea, we targeted the structure of CB2R bound to the inverse agonist, AM10257 (PDB 5ZTY)^8^. Here again, we optimized the sampling to engage with polar residues T3.33, S7.39, S2.60, H.265 and the backbone of L182^ECL2^ and calculated energy potential grids that, in control calculations with property-matched decoys, led to favorable enrichments (**SI Figure S9**). Returning to the higher sampling of the first, in-stock library campaign, we docked a library of 2.6 billion make-on-demand molecules against this inactive state, sampling each in an average of 579,700 configurations. The molecules were filtered using the same criteria as before, and also for distances nearing toggle switch residue, W6.48 (Figure 2A) to promote stabilization of the inactive state^50^. This led to 75,003 molecules with docking scores better than −25. Clustering for diversity led to 4,839 sets. These were visually inspected and 327 were prioritized for make-on-demand synthesis at Enamine, of which 290 were successfully made (a fulfillment rate of 89%).

**Figure 2.**
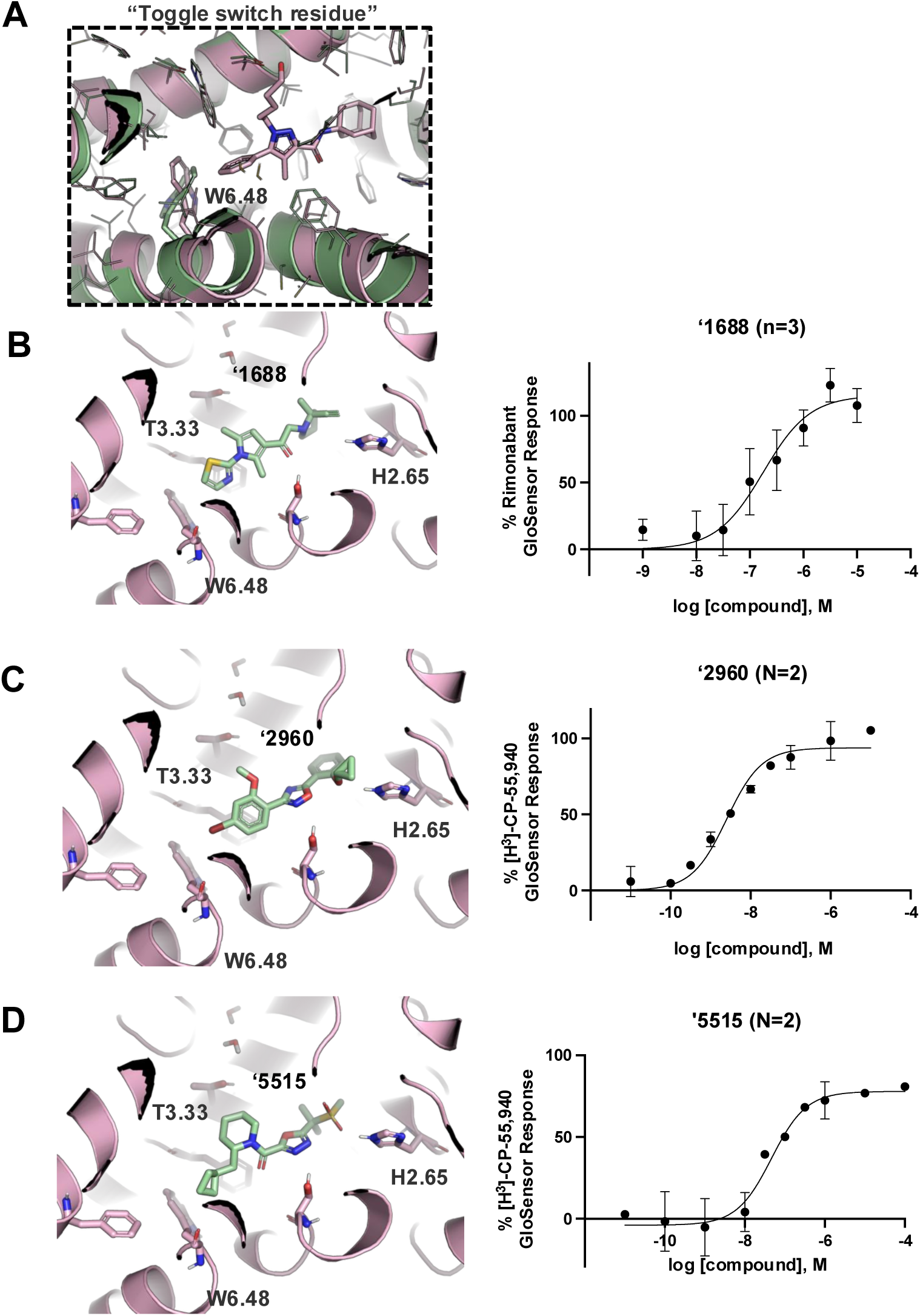
Active (green) and inactive (pink) structures. Distances to W6.58 was an additional filter for the 2.6B screen against the inverse agonist bound structure to stabilize the inactive conformation (**A**). Representative inverse agonists (**B**) and agonists (**C** and **D**) docked into the inactive state, with concentration response curves. Inverse agonism was determined by monitoring cAMP accumulation with GloSensor in the presence of Forskolin and CP-55,940 normalized to Rimonabant. Agonism was determined by monitoring cAMP accumulation with GloSensor in the presence of Forskolin normalized to CP-55,940. n=technical repeats, N=independent experiments.

These 290 molecules were initially tested at 10 μM by radio-ligand displacement. Of these, 89 showed ≥ 30% displacement (**Supporting Information 2.6B_Screen.xlsx**) and 69 having ≥ 40% displacement were advanced into full concentration response. Of these, 49 had K_i_ values ≤15 µM by Cheng-Prusoff^43^ (SI **Figure S10**), a hit rate of 17%, while 10 had K_i_ values ≤1 μM. The 10 best compounds (**Table 3**) were followed up with concentration response curves using cAMP levels as a readout (Glosensor) in the presence of dual CB1R/CB2R agonist CP55,940. Three were inverse-agonists the best of which had an EC_50_ of 28 nM, while five were agonists the best of which had an EC_50_ 2.5 nM with two others in the same range (Figure 2B, **SI Figure S11**).

**Table 3.**
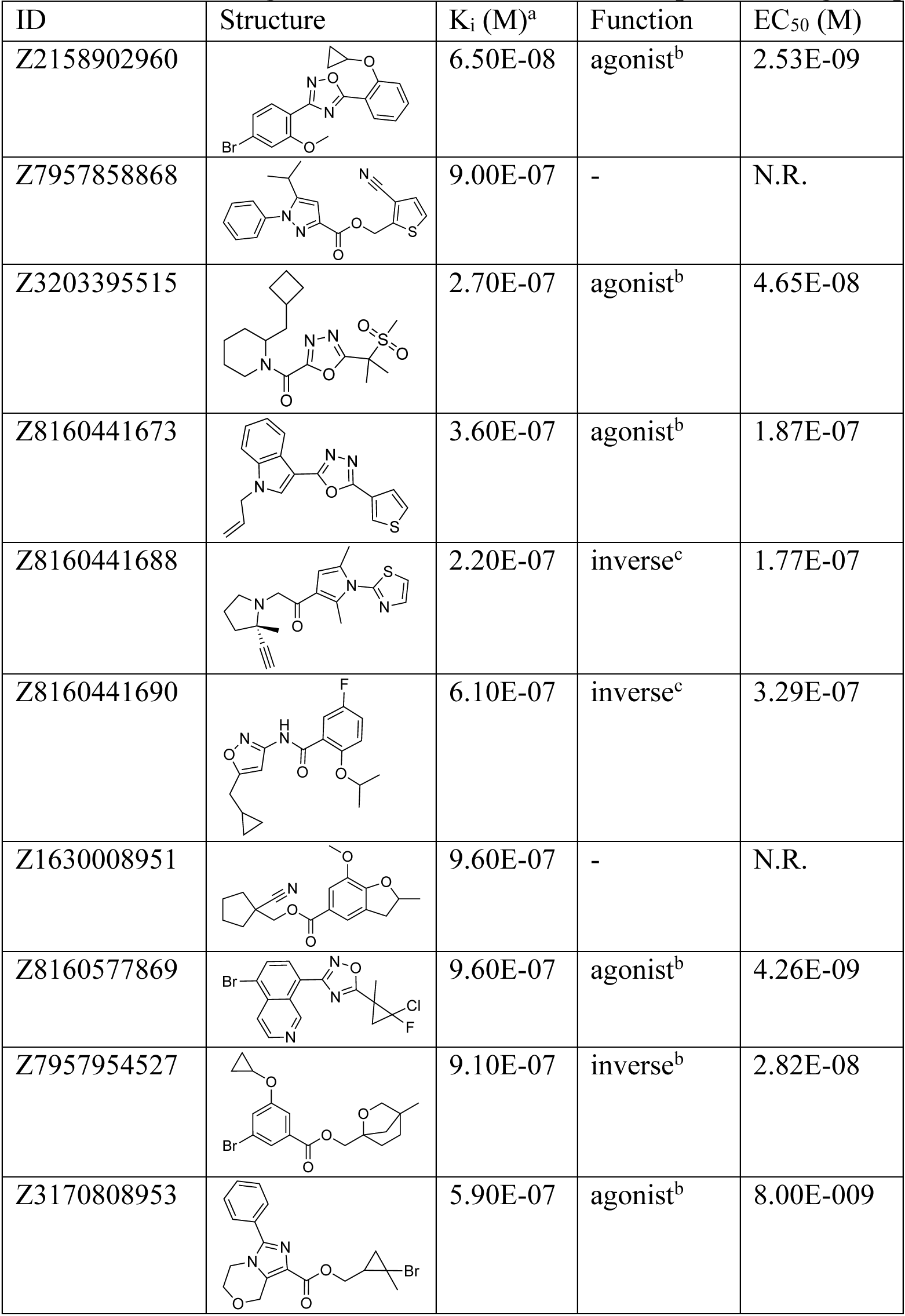
Ten CB2R ligands from the 2.6B inactive receptor docking campaign. **a.** K_i_ values determined from displacement of [^3^H]-WIN 55212-2 minus non-specific binding. **b.** Agonism was determined by monitoring cAMP accumulation with GloSensor in the presence of Forskolin normalized to CP-55,940. N.R. = No Response. **c.** Inverse agonism was determined by monitoring cAMP accumulation with GloSensor in the presence of Forskolin and CP-55,940 normalized to Rimonabant.

| ID | Structure | K<sub>i</sub> (M)<sup>a</sup> | Function | EC<sub>50</sub> (M) |
| --- | --- | --- | --- | --- |
| Z2158902960 |  | 6.50E-08 | agonist<sup>b</sup> | 2.53E-09 |
| Z7957858868 |  | 9.00E-07 | - | N.R. |
| Z3203395515 |  | 2.70E-07 | agonist<sup>b</sup> | 4.65E-08 |
| Z8160441673 |  | 3.60E-07 | agonist<sup>b</sup> | 1.87E-07 |
| Z8160441688 |  | 2.20E-07 | inverse<sup>c</sup> | 1.77E-07 |
| Z8160441690 |  | 6.10E-07 | inverse<sup>c</sup> | 3.29E-07 |
| Z1630008951 |  | 9.60E-07 | - | N.R. |
| Z8160577869 |  | 9.60E-07 | agonist<sup>b</sup> | 4.26E-09 |
| Z7957954527 |  | 9.10E-07 | inverse<sup>b</sup> | 2.82E-08 |
| Z3170808953 |  | 5.90E-07 | agonist<sup>b</sup> | 8.00E-009 |

These results are broadly consistent with earlier studies that found little ability to select for agonists or inverse agonists by targeting different receptor states. While the screen against the active state did appear to show a bias toward agonists, this may reflect the small number of compounds tested, and their micromolar activities. The campaign against the inactive state, with many more compounds tested and far more potent compounds found, may be the more informative and it suggests that docking against an inactive state, at least, little-biased the actives to inverse-agonists. Indeed, the most potent molecules to emerge from this screen were agonists.

That said, the potency of the molecules that did emerge was high relative to the 7 million molecule library campaign that had similar sampling, with the best inverse agonist from the billion molecule campaign 4-fold more potent than the most active molecule from the smaller library, and the best agonist over 14-fold more potent by binding. Importantly, no functional activity could be measured from the smaller library screen, whereas mid- and low-nM range inverse agonists and agonists were found from the large library campaign. Meanwhile, the new actives were topologically unrelated not only to previously known ligands, but also the molecules discovered from the two campaigns against the active state structures, points that we consider below.

### Hit rates and affinities from the larger and smaller library docking versus CB2R

We have previously found that as libraries grow docking scores improve, and improved scores lead to improved hit rates and affinities^31, 51–54^. In this study, too, there were about 200-fold more high-scoring molecules across the spectrum in the 2.6 billion molecule library screen versus that of 7 million molecule library screen, and the scores left-shifted to more negative (better) scores, reflecting the increase library size (Figure 3A). Experimentally, these better scores were reflected in more potent ligands discovered from the larger library (Figure 3B).

**Figure 3.**
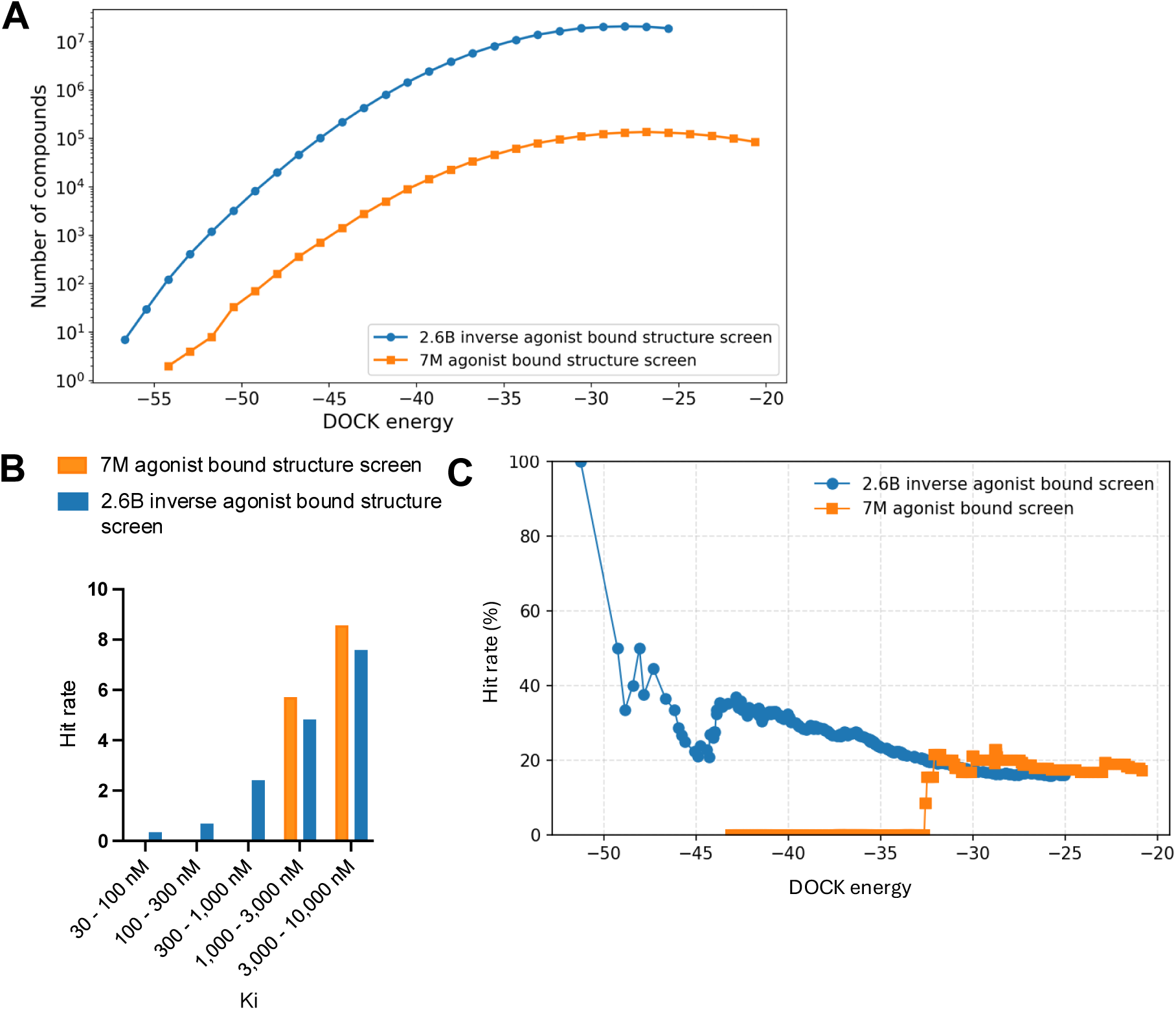
Docking scores improve in the 2.6 billion molecule screen versus the 7 million molecule screen (**A**). The K_i_ distribution from the 2.6 billion molecule and the 7 million molecule docking campaign (**B**). The correlation between docking scores and hit rates for the 2.6 billion and 7 million molecule library screens (**C**).

The K_i_ values of the ligands improved substantially from the larger versus the smaller library docking campaigns, and the difference are more exaggerated still comparing functional activities, as none of the docking hits from the smaller library campaign had measurable functional activity as agonists or inverse agonists whereas from the large library agonists and inverse agonists active in the mid- and low-nM range were found. As in earlier studies, hit rates climbed as docking scores improved, and the larger library achieved higher hit rates in the higher scoring range (Figure 3C). A caveat is that the smaller library campaign, with only 34 molecules tested experimentally, was underpowered.

### Ligand optimization

If one way to find more potent ligands is to dock larger libraries, a second is to optimize initial actives, something that has the added advantage of investigating the trustworthiness of one’s hits, as optimizability within an understandable SAR typically means that a series is mechanistically trustworthy. This was a concern for the small library actives which, while they had well-behaved ligand-displacement curves, were only µM in affinity and had no detectable functions as agonists or inverse agonists (which is not uncommon for early GPCR actives^38, 51^). Accordingly, we picked exemplar compounds from the high-sampling 7 million molecule and the lower-sampling 1.6 billion molecule campaigns and attempted to optimize them for affinity (Figure 4, **SI Figure S12**).

**Figure 4.**
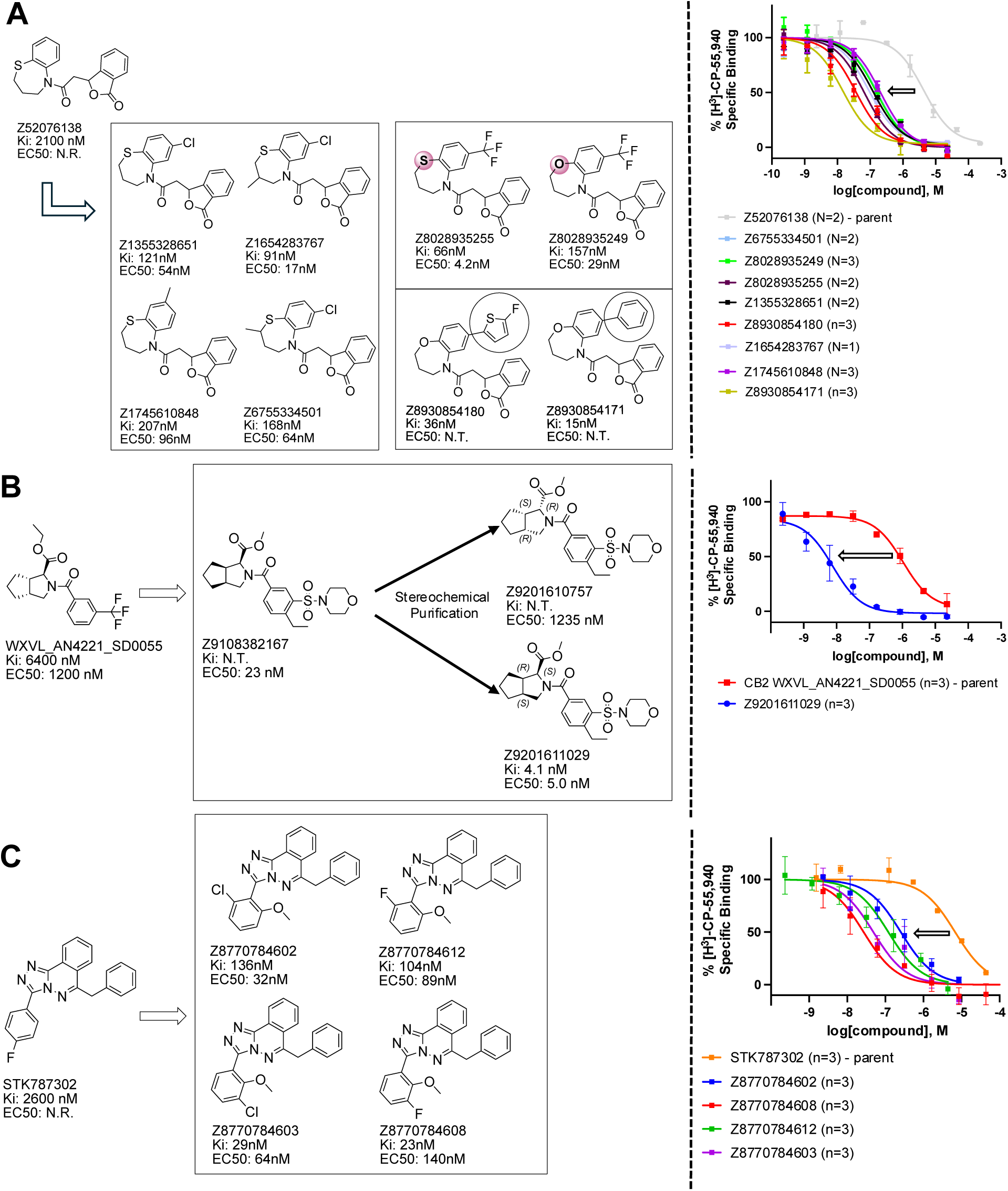
Analogs ≤100 nM of ‘6138 from the 7M screen (**A**), ‘0055 from the 1.6B screen (**B**) and ‘7302 from the 7M screen (**C**) along with their binding curves. K_i_ values were determined from displacement of [^3^H]-CP-55,940 minus non-specific binding. Agonism was determined from BRET signals monitoring G_i_ activation normalized to Fubinaca signals.

The eight best analogs of ‘6138 (2.1 μM) from the 7 million library screen showed 10- to 140-fold improvement in Ki. Whereas the parent had no measurable activity in BRET functional assays, the analogs had functional EC_50_‘s in the low- to mid-nM ranges (Figure 4A, **SI Figure S12A**). Small modifications were readily available from the billion-molecule make-on-demand set and could be accommodated by the binding site based on docking poses (Figure 5A). Further optimization included increasing polarity from ‘5255 (cLogP 3.9) to ‘5249 (cLogP 3.2), which led to 115-fold selectivity for CB2R over CB1R for the analog ‘5249, while ‘5255 was 51x more selective (**SI Figure S13**).

**Figure 5.**
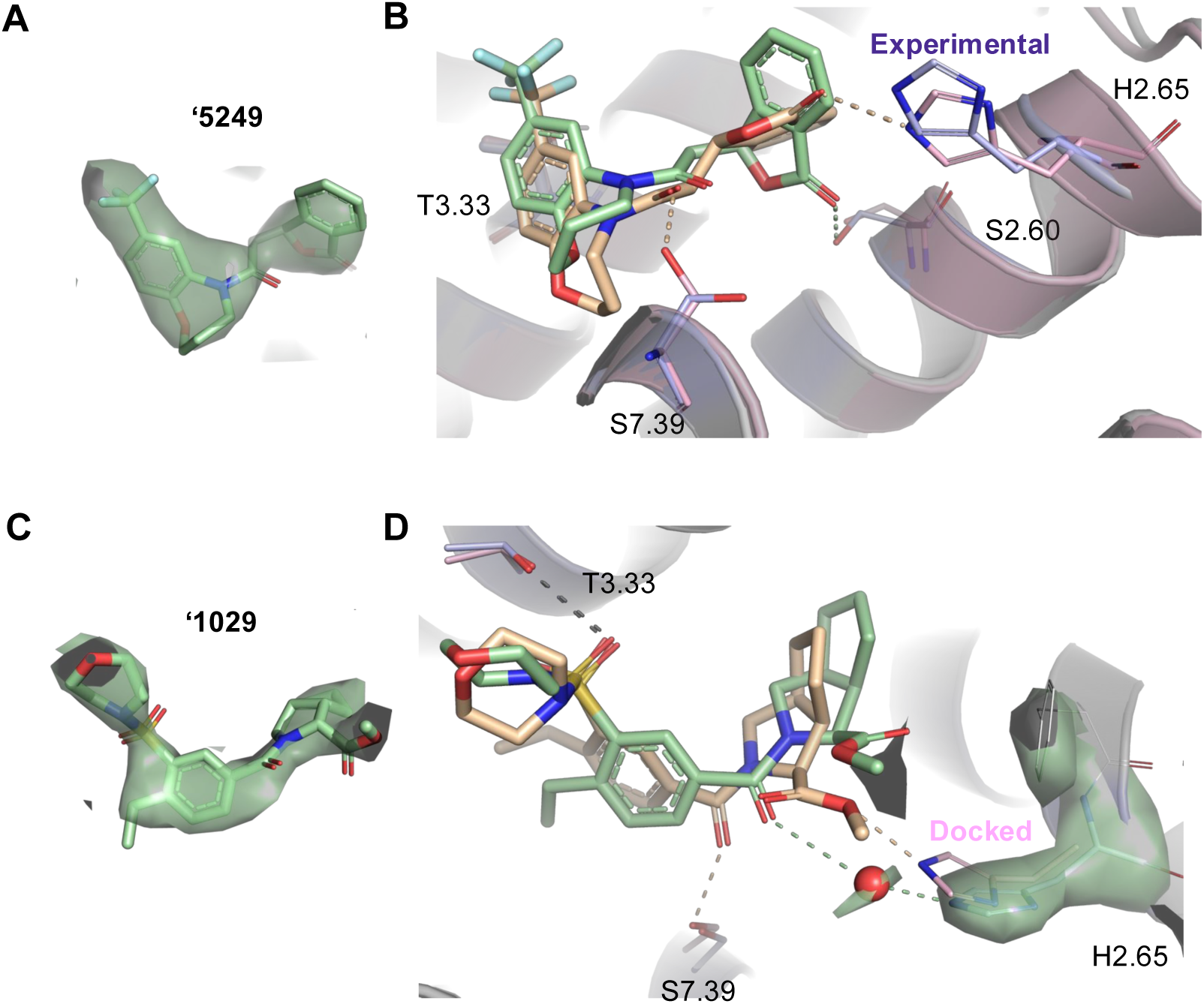
Cryo-EM structures of two of the new CB2 receptor ligands. Cryo-EM density of ‘5429, an optimized analog deriving from the 7 million molecule library screen (**A**). Superposition of the docking (tan) and experimental (green) poses of ‘5429 in the CB2R orthosteric site (**B**). Predicted and observed hydrogen-bonds are shown as dashed lines. Cryo-EM density of ‘1029, an optimized analog deriving from the 1.6 billion molecule library screen (**C**). Superposition of the docking (tan) and experimental (green) poses of ‘1029 in the CB2R orthosteric site (**D**). The densities in the figure are shown as surfaces contoured at a threshold of 20 for ‘2167 and 17 for ‘1029. An overall resolution of 2.9Å was obtained for both structures.

The best analog of ‘0055 from the 1.6B screen (EC_50_ 1.2 μM) improved activity 50-fold (Figure 4B), modeled to occur via interactions in a CB2R sub-pocket near T3.33, which the newly installed sulfonyl was also predicted to engage through a hydrogen-bond (Figure 5B). The conversion of the stereochemistry around the fused pyrrolidine ring from trans- to cis-also seems to have contributed to the better affinity, and stereochemical purification of ‘2167 led to an EC_50_ of 5.0 nM for the cis isomer, ‘1029 (**SI Figure S14**). Finally, systematic small perturbations of ‘7302 (Ki: 2.6 μM) from the 7 million library screen resulted in up to 118-fold improvements in affinity. These studies suggest that the initial docking hits were bona fide ligands and provide potent members for each of the three scaffold classes.

### Cryo-EM structures

To complement the structure-based optimization, guide future work, and to test the docking predictions, cryo-EM structures of ligand-bound CB2R were determined for agonists ‘5249 and ‘1029, each representing a different scaffold family. For both agonists, the docked and the experimental pose superposed well (Figure 5) with ligand RMSD’s of 1.8 and 1.1 Å, respectively. Intriguingly, the experimental structure of ‘5249 revealed an S instead of R isomer, although the racemic mixture showed similar activity values (**SI Figure S15**). Due to this, the predicted hydrogen-bond with S7.39 and H2.65 did not occur while instead a hydrogen-bond with S2.60 was observed. For ‘1029 the correct isomer was predicted, however, the interaction with S7.39 and H2.65 was again lost. Instead, an observed EM density points towards a possible water mediated interaction with H2.65. Both the experimental and predicted models retain an apparent hydrogen-bond with T3.33.

### The diversity and novelty of the new ligands

Only molecules that were topologically unrelated to known cannabinoids in ChEMBL were selected from the docking. Overall, 71 diverse ligands were discovered with K_i_ values ≤15 µM, and 16, discovered via docking and optimization (**Table 3**, Figure 4), had activities ≤100 nM. These 16 could be clustered into 8 ligand families using ECFP4 Tc values ≤0.35 to any other family as a clustering criterion^55^. Comparing the compound in each of these eight families with the highest Tc to the 2,391 ≤100 nM CB2R ChEMBL ligands reveals that most have little similarity to any previously known family of CB2R ligands (Figure 6). Only chemotype 3 resembles a known potent CB2R ligand (Tc 0.48). This is an example of analoging drifting into a known chemotypes, as the parent docking hit in chemotype 3 was dissimilar to the known ligands (Tc ≤0.35). For four families where we measured selectivity for CB2R versus the highly-related CB1R, we found agonists that were between 8- and 115-fold selective. In chemotype family 1, the most selective from those tested was ‘5249 (Figure 4), having 115-fold selectivity for CB2R (**SI Figure S13, SI Figure S16A**). From chemotype family 2, compound ‘4602 was 45-fold more selective (**SI Figure S16B**). From chemotype family 3, ‘1029 was 8-fold selective for CB2R over CB1R (**SI Figure S16C**), while compound ‘2960 from chemotype family 4 was 14-fold selective for CB2R over CB1R (**SI Figure S16D**).

**Figure 6.**
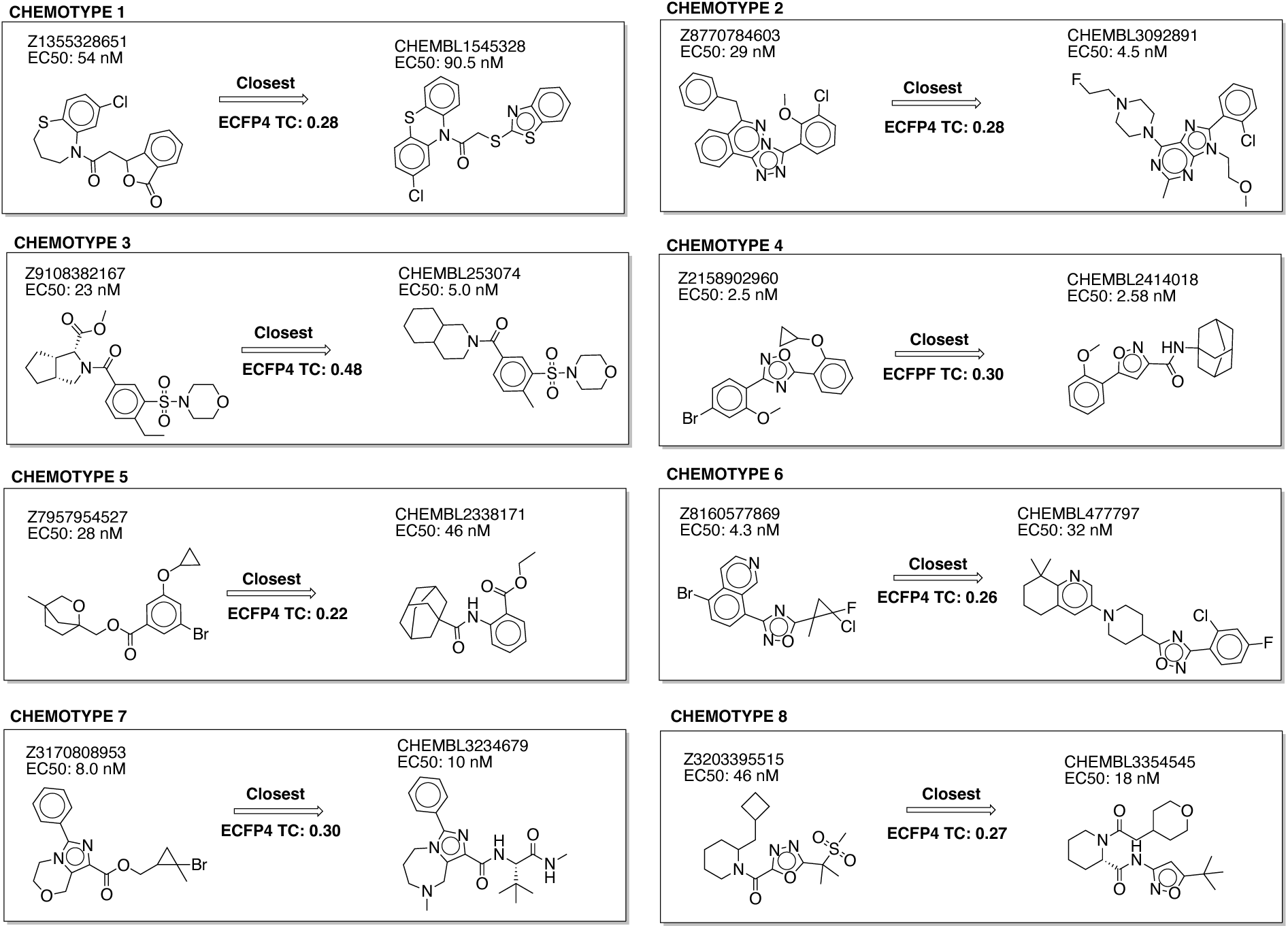
The eight chemotypes discovered in this work are diverse (ECFP4 Tc values ≤0.35). Seven out of the eight are novel (ECFP4 Tc values ≤0.35 versus 2,391 known cannabinoid ligands ≤100 nM in ChEMBL). RDKit was used for clustering by similarity and novelty calculations.

## DISCUSSION AND CONCLUSIONS

Four observations from this study merit emphasis. **First**, even in relatively non-polar site as a lipid-receptor like CB2R, polar interactions may be found and biasing toward them appears to lead to more selective ligands. **Second,** docking hit rates, affinities, and functional effects improved as we moved from smaller to larger libraries. Whereas docking an in-stock set of 7 million molecules led to ligands in the µM range, those from a 2.6 billion molecule library had at least 14-fold better affinities and the difference was far greater still for functional activities, which reached low nM values from the large library but were not detectable from the small library docking screen. As docking scores improved, hit rate rose. This has been seen previously against unrelated targets, and hints at a loose association between docking score and not only hit rate but perhaps also affinity^31^, arguably supporting further efforts to grow docking libraries into the hundred-billion and trillion-molecule make-on-demand spaces, and potentially even larger spaces^56, 57^ now available. A way to access these libraries may be via synthon methods^19, 58, 59^. Supporting the ability of such synthon approaches, brute-force docking of a 2.6 billion molecule library found similar hit rates as synthon docking against CB2R that sampled from an 11.5 billion molecule library, with the synthon method explicitly docked far fewer molecules. **Third,** although more potent ligands emerged from docking the 2.6 billion molecule library against the inactive conformation of CB2R, versus the 7 million molecule library, docking against the inactivated state did not strongly bias toward inverse agonists over agonists—potent examples of each were found. This is consistent with our experience against other GPCRs^38, 47, 51, 54, 60^, with only a few exceptions^61, 62^. It may also be noted that CB2R active and inactive state pocket residues differ <u><1.5Å compared to other GPRC’s for example CB1R where >2.5Å differences are observed,</u> hampering the distinction between agonists and inverse agonists through structure-based approaches for this target. As such, docking with fidelity to functions encoded in ligand-bound GPCR structures remains an unmet goal in the field. **Finally,** this study revealed three new families of CB2R agonists with EC_50_‘s ≤10 nM and four more with activities between 29 to 54 nM, with chemotypes unrelated to cannabinoid ligands in ChEMBL^32^. These molecules are 8 to 115-fold selective for CB2R versus CB1R. Supported by the cryoEM structures determined here, they may template new probe discovery efforts against this fascinating receptor.

Several caveats merit airing. Our confidence in the 7 million molecule in-stock set docking are affected by the relatively few molecules tested. Whereas testing around 35 docking hits remains common in the field, this small a number does lead to uncertainties around hit rates and affinity ranges^31^. Our comparison of the 2.6 billion library screen to the smaller in-stock set is also clouded by the active-state and inactive state receptor structures, respectively, that were targeted. While the hit rates were similar between the full docking of the 2.6 billion molecule library and the implied docking of 11.5 billion molecules in a synthon-based method^19^, the affinities of the actives from the full (brute-force) docking of the 2.6 billion molecule library were 4-fold more potent in binding assays and 44-fold more potent in functional assays. Given that the synthon docking explicitly treated 10^4^-fewer compounds, this drop-off is not unexpected. One caveat here is that far more molecules were selected for testing. While docking scores, hit rates, and affinities improved with the larger versus the smaller library, any correlation between docking scores, affinities and hit rates remained very weak; how to improve this correlation remains stubbornly resistant to what should be improvements in scoring^63, 64^ and remains an active area of research in the field, including through ongoing docking-and-testing competitions^65, 66^.

These caveats should not obscure the central observations for this study. Even in lipid receptor sites, prioritizing polar complementarity can lead to better physical properties and improved sub-type selectivity among docking-derived ligands. In CB2R, as in other targets^31, 54^ interrogating larger libraries revealed more potent actives than did smaller libraries, supporting ongoing efforts to further grow molecular libraries for virtual screening. From these studies, a broad spectrum of chemically distinct and potent CB2 receptor agonists and inverse agonists emerged.

## EXPERIMENTAL SECTION

### Docking model optimization and virtual screening

The CB2 PDB structure, 6PT0^30^, bound to agonist WIN 55,212-2 was used for the 7M and 1.6B screens, and the CB2 PDB structure, 5ZTY^8^, bound to inverse agonist AM10257 was used for the 2.6B screen. Both structures were protonated using Schrödinger’s Maestro Preparation Wizard^67^. Rotamers^68^ and protonation states of polar pocket residues (T3.33, S7.39, S2.60, H.265 and L182^ECL2^) were optimized within density for increased hydrogen-bond interaction potential.

The coordinates of cryogenic ligand atoms were used for placing pre-generated molecule conformers from the ZINC22 database (https://zinc22.docking.org) into the binding site. Additional coordinates were placed appropriately when the ligands could not interact with the polar pocket residues mentioned above. A total of 45 experimentally derived, manually defined and randomly assigned sampling coordinates were used for all screens. The three parameters “match goal” (the number of orientations per molecule allowed to be sampled), “bump_rigid” and “bump_maximum” (steric filters during orientational sampling) for the 7M and 2.6B screens were set to 2000, 20 and 20, while these parameters were reduced to 1000, 10 and 10 for the 1.6B screen.

This led to a sampling of about 1700 orientations per molecule for the 7M and 2.6B screens, while the 1.6B screen gave about 450 orientations per molecule. About 330 conformers per orientation were sampled for each screen.

To rank the sampled orientations by complementarity to the binding pocket, grids were pre-calculated using CHEMGRID^69^ for AMBER^70^ van der Waals potentials, QNIFFT^71^ for Poisson-Boltzmann-based electrostatic potentials, and SOLVMAP^72^ for context-dependent ligand desolvation. Retrospective studies using 34 well-characterized CB2 ligands extracted from the IUPHAR database^34^ and 1690 property-matched but topologically dissimilar molecules (decoys) from DUDE-Z^35^ were used to optimize docking parameters based on the ability to prioritize ligands over non-ligands (measured by logAUC) and geometric fidelity of molecules with experimentally derived binding modes.

Optimization included defining the extended low protein dielectric and desolvation region^73^, using SPHGEN^74^. The protein low dielectric and desolvation regions were extended, based on control calculations, by a radius of 1.0 Å and 0.1 Å for the agonist bound model, and 1.3 Å and 0.2 Å for the inverse agonist bound model. Furthermore, electrostatic grid potentials corresponding to polar heteroatoms of residues T3.33, S7.39, S2.60, and the backbone of L182^ECL2^ were empirically scaled by −0.4 grid units, with compensatory scaling of polar hydrogen grid potentials by +0.4, to enhance discrimination of complementary polar interactions. As a final sanity check, for the agonist bound structure, we ensured prioritization of neutral over charged molecules (**SI Figure S17**).

The 7M, 1.6B and 2.6B libraries comprised neutral molecules that were less than 29 heavy atoms with cLogP ≤5. A DOCK3.8 score filter of −20, −25 and −25 was applied, which led to 1.4M, 22M and 164M molecules. Filtering with LUNA^75^ for hydrogen-bonds with T3.33, S7.39, S2.60, H.265 or L182^ECL2^, and excluding molecules with a hydrogen-bond donor that lacked a complementary acceptor in the orthosteric site, and that had >3 hydrogen-bond acceptors without a complementary receptor donor, left 101K, 146K and 75K molecules (including an additional filter removing molecules with distances >4Å to W6.48 for the 2.6B screen). Filtering for novelty (ECFP4 0.35 Tc against 341 cannabinoid receptor ligands with ECFP4 Tc values ≤0.35 amongst each other extracted from ChEMBL at a K_i_ cutoff of 10 μM), and clustering for diversity (ECFP4 Tc 0.35), left about 7K, 10K and 5K molecules. The top-ranking molecules were manually inspected for favorable geometries and diversified interactions.

### Hit optimization

Analogs of hits were sought through a combination of similarity and sub-structure searches of the Arthor (https://arthor.docking.org) and SmallWorld (https://sw.docking.org/) databases from a 46-billion- molecule make-on-demand library, or were designed based on specific hypotheses synthesizable at Enamine. Potential analogs were docked to either the agonist or inverse agonist bound models. Docked analogs that maintained geometric overlap with the predicted pose of the parent molecule were filtered using the same criteria a above. After visual inspection, prioritized analogs were acquired and tested experimentally.

### Synthesis of molecules

The in-stock prioritized molecules were sourced from Enamine, MCule and MolPort. From the 1.6B screen, 151 molecules prioritized for purchasing were synthesized by Enamine (sourced from https://enamine.net/compound-collections/real-compounds) and WuXi (sourced from https://chemistry.wuxiapptec.com/small_molecule) with a fulfillment rate of 79%. From the 2.6B screen, all molecules were synthesized by Enamine with a fulfillment rate of 89%. The purities of active molecules were at least 90% and typically greater than 95%. Synthetic routes, chemical characterization, and purity information of hits and analogs can be found in Supporting Information Synthetic_procedures.docx and Spectra.docx.

### CB1/CB2 competition binding assays for the 7M and 1.6B screens

The binding affinities of the compounds were obtained by competition binding using membrane preparations from HEK293 cells stably expressing human CB1 or CB2 receptors and [^3^H]-CP-55,940 as the radioligand, as previously described^9,22^. The results were analyzed using nonlinear regression to determine the IC_50_ and K_i_ values for each ligand (Prism by GraphPad Software, Inc., San Diego, CA).

### CB2 competition binding assays for compounds from the 2.6B screen

The binding affinities of the compounds were obtained by competition binding using membrane preparations from CHO cells stably expressing human CB2 receptors and [^3^H]- WIN 55212-2 as the radioligand, as previously described^76^. The results were analyzed using nonlinear regression to determine the IC_50_ and K_i_ values for each ligand. This analysis was performed using software developed at Cerep (Hill software) and validated by comparison with data generated by the commercial software SigmaPlot® 4.0 for Windows® (© 1997 by SPSS Inc.).

### Dynamic Light Scattering (DLS)

Similar to what has been previously described^77^, samples were prepared in filtered 50 mM KPi buffer, pH 7.0 with final DMSO concentration at 1% (v/v). Colloidal particle formation was detected using DynaPro Plate Reader II (Wyatt Technologies).

All compounds were screened in triplicate at 100μM and, if colloids were detected, 8-point half-log dilutions of compounds were performed on DLS in triplicate. To determine the critical aggregation concentration (CAC), data for each compound were spilt into two data sets based on aggregating (i.e. >10^6^ scattering intensity) and non- aggregating (i.e. <10^6^ scattering intensity) scattering intensities and were fitted with separate nonlinear regression curves. The point of intersection was determined using GraphPad Prism software version 10.1.1 (San Diego, CA).

### Enzyme Inhibition Assays

Enzyme inhibition assays against counter-screen enzymes were performed at room temperature using CLARIOstar Plate Reader (BMG Labtech). Samples were prepared in 50 mM KPi buffer, pH 7.0 with final DMSO concentration at 1% (v/v). Compounds were incubated with 2 nM AmpC β-lactamase (AmpC) purified as previously described^78^ or malate dehydrogenase (MDH) (Sigma-Aldrich, 442610) for 5 min. AmpC reactions were initiated by the addition of a 50 μM CENTA chromogenic substrate (Sigma-Aldrich, 219475). The change in absorbance was monitored at 405 nm for 80 s. MDH reactions were initiated by the addition of 200 μM nicotinamide adenine dinucleotide (NADH) (Sigma-Aldrich, 54839) and 200 μM oxaloacetic acid (Sigma-Aldrich, 324427). The change in absorbance was monitored at 340 nm, also for 80 s. Initial rates were divided by the DMSO control rate to determine the percent enzyme activity (%). Each compound was initially screened at 100 μM in triplicate against MDH. If no inhibition was observed against MDH, compounds were screened against other counter-screen enzyme AmpC at 100 μM. If a given compound showed ≥30% inhibition against either counter-screen enzyme, a concentration-response curve was performed in triplicate to determine IC_50_. Data was analyzed using GraphPad Prism software version 10.1.1 (San Diego, CA).

For detergent reversibility experiments, compounds that showed ≥ 30% enzyme inhibition against either MDH or AmpC were screened at 100 μM with or without 0.01% (v/v) Triton X-100 in triplicate. Enzymatic reactions were performed/monitored as described above.

### Plasmids and Baculovirus Generation

Human CB2 bearing an N-terminal hemagglutinin signal peptide-FLAG epitope tag and dominant-negative human Gαi1 (DNGi1) containing the mutations S47N, G203A, E245A, and A326S, were subcloned into individual pFastBac1vectors. Human Gβ1, engineered with an N-terminal 8×His tag, and bovine Gγ2 were subcloned into the pFastBac-Dual vector (Invitrogen) under polyhedrin and p10 promoters, respectively. Recombinant bacmids were generated via Tn7-mediated transposition in DH10Bac *Escherichia coli* cells using the Bac-to-Bac system. Baculovirus was produced and amplified in *Spodoptera frugiperda* (Sf9) insect cells over three passages (∼12–14 days).

For G protein signaling assays, human CB2 and CB1 containing an N-terminal hemagglutinin signal peptide and FLAG epitope tag were cloned into pcDNA3.1. Plasmids encoding the Gαi1-RLuc8, Gβ, and Gγ-GFP2 biosensors were obtained from the TRUPATH platform^79^.

### Bioluminescence Resonance Energy Transfer (BRET) Signaling Assays

CB2- and CB1-mediated G protein signaling was measured using the TRUPATH BRET assay, which reports agonist-induced dissociation of Gα-RLuc8 from Gβγ-GFP2. Suspension HEK293 cells were cultured in FreeStyle 293 Expression Medium (Thermo Fisher Scientific) to a density of ∼1.25 × 10⁶ cells/mL and transiently transfected with an equimolar mixture of CB2 or CB1, Gα-RLuc8, Gβ, and Gγ-GFP2 plasmids using polyethylenimine (PEI). Transfections were performed with 600 ng total DNA and 1,800 ng PEI per mL of culture, corresponding to a 1:3 DNA:PEI mass ratio.

After 48–72 h of expression, cells were harvested by centrifugation (700 × g, 1 min) and gently resuspended in assay buffer consisting of 1× Hank’s balanced salt solution (HBSS), 0.1% bovine serum albumin (BSA), and 6 mM MgCl₂. Serial dilutions of test compounds were prepared in assay buffer and 30 µL transferred to 96-well opaque plates (Corning). The CB2/CB1 full agonist AB-fubinaca was included on each plate as a positive control. Cells were resuspended in HBSS containing 5 µg/mL coelenterazine 400a (Cayman Chemical) and added to assay plates to a final volume of 90 µL per well.

BRET signals were acquired at room temperature over a 15 min period using a SpectraMax iD5 plate reader with GFP2 filter settings, monitoring donor and acceptor emission wavelengths of 410 nm and 515 nm. Computed BRET ratios (GFP2 or mVenus/RLuc8) at 00:03:50, 00:07:36, and 00:11:23 (hh:mm:ss) were averaged, and ligand-free well values were subtracted from ligand-treated well values to obtain net BRET signals. Net BRET values were normalized to the AB-fubinaca E_max_ and plotted as a function of ligand concentration. Data from biological triplicates were fit using a three-parameter logistic nonlinear regression model in GraphPad Prism.

### GloSensor cAMP accumulation assay

The GloSensor cAMP assay for human CB2 was performed following the manufacturer’s instructions (Promega) with minor modifications. Briefly, WT human CB2 was cloned into pcDNA3.1 and co-transfected with the 22F cAMP GloSensor plasmid (Promega) into HEK293T cells cultured in 6-well plates. At 24 h post-transfection, cells were dissociated and reseeded into white, clear-bottom 96-well plates in CO₂-independent medium (Thermo Fisher) supplemented with the GloSensor cAMP reagent (diluted according to the manufacturer’s instructions). Plates were then incubated for 1 h at 37 °C followed by 1 h at room temperature in the dark to allow equilibration of the reporter. To elevate basal cAMP for Gi/o-coupled CB2 readout, 10 µM forskolin was added to all wells immediately prior to compound addition. Serial dilutions of test ligands (or vehicle) were then added, and luminescence was measured on a microplate luminometer. Data were normalized to the forskolin-only condition (100%) and fitted in GraphPad Prism 9.0 to obtain EC₅₀/IC₅₀ values for inhibition of cAMP. Where indicated, reference agonists (CP55,940) were included as positive controls, and CB2 inverse agonists were evaluated by 10–15 min preincubation with 10 nM CP55,940 before agonist addition.

### Expression and Purification of the CB2–Gi1 Complex

Human CB2 was co-expressed with dominant-negative Gαi1 (DNGi1) and Gβγ in *Spodoptera frugiperda* (Sf9) insect cells using the baculovirus expression system. Sf9 cultures (6 L total volume) were grown to a density of 3.0 × 10⁶ cells/mL and co-infected at a multiplicity of infection of 1:1:1 for CB2, DNGi1, and Gβγ. Cells were harvested 72 h post-infection at approximately 70% viability, pelleted by centrifugation, and stored at −80 °C until use.

Frozen cell pellets were thawed and resuspended in lysis buffer containing 20 mM HEPES (pH 7.5), 100 mM NaCl, 10 mM MgCl₂, 100 µM TCEP, 25 mU/mL apyrase (New England Biolabs), 10 µM CB2 agonist, and protease inhibitors (1 mM benzamidine, 100 µg/mL leupeptin, 2 µg/mL aprotinin, and 2 µg/mL pepstatin). Cells were disrupted by nitrogen cavitation (750 psi, 30 min, 4°C) using a Parr cell disruption bomb, and lysates were clarified by ultracentrifugation at 100,000 × g for 30 min.

Membrane pellets were resuspended in lysis buffer and solubilized for 3 h at 4 °C with gentle stirring in the presence of 1% lauryl maltose neopentyl glycol (LMNG) and 0.2% cholesteryl hemisuccinate (CHS). Insoluble material was removed by ultracentrifugation, and the solubilized fraction was supplemented with 30 mM imidazole prior to incubation with Ni–NTA resin for 1 h at 4 °C. The resin was washed with buffer containing decreasing concentrations of LMNG/CHS and eluted with 250 mM imidazole.

The eluate was supplemented with 2 mM CaCl₂ and applied to anti-FLAG M1 resin equilibrated in FLAG wash buffer (20 mM HEPES pH 7.5, 100 mM NaCl, 5 mM CaCl₂, 0.01% LMNG/0.001% CHS, 10 µM agonist). After washing with ∼5 column volumes, the complex was eluted in buffer without CaCl₂ and containing 0.2 mg/mL FLAG peptide and 2 mM EDTA.

Eluted protein was concentrated using a 100 kDa molecular weight cutoff centrifugal concentrator and further purified by size-exclusion chromatography on an ENrich SEC 650 column (Bio-Rad) equilibrated in buffer containing 20 mM HEPES pH 7.5, 100 mM NaCl, 0.001% LMNG, 0.00033% glyco-dioleyl-neopentyl glycol (GDN), 0.0001% CHS, and 20 µM agonist. The CB2–Gi1 complex eluted as a single monodisperse peak at ∼12.25 mL and was concentrated to ∼5–10 mg/mL for cryo-EM analysis.

### Cryo-Electron Microscopy Grid Preparation, Data Collection, and Processing

Purified CB2–Gi1 complex (3 µL at 5–10 mg/mL) was applied to glow-discharged UltrAuFoil R1.2/1.3 gold grids (Quantifoil). Grids were blotted for 3 s at 4 °C and 100% relative humidity using a Vitrobot Mark IV (Thermo Fisher Scientific) and plunge-frozen into liquid ethane.

Cryo-EM data were collected on a Titan Krios G2 transmission electron microscope operated at 300 kV and equipped with a Falcon 4i direct electron detector and Selectris energy filter (Thermo Fisher Scientific). Data were acquired using EPU software at a calibrated pixel size of 0.96 Å, with a cumulative electron dose of 50 e⁻/Å² and a defocus range of −0.4 to −1.6 µm.

Cryo-EM data processing was performed using CryoSPARC (Structura Biotechnology). Movies were subjected to patch-based motion correction and contrast transfer function estimation prior to particle extraction. Particles were iteratively filtered through multiple rounds of two-dimensional classification and refined using successive rounds of *ab initio* reconstruction and heterogeneous refinement. A final subset of particles was used for nonuniform refinement, yielding reconstructions at an overall resolution of 2.9 Å for both structures based on the gold-standard Fourier shell correlation criterion (FSC = 0.143). Resolution of the CB2 receptor region was further improved through local refinement.

## ASSOCIATED CONTENT

### Supporting Information

- Figures showing optimization of retrospective docking, Results of colloidal aggregation assays, Binding curves of initial docking hits, Functional activity and selectivity of docking hits, Activity and selectivity of (purified) analogs (Supporting-Information.docx)
- ID’s, SMILES, Docking scores, % displacement and K_i_ values of tested compounds from the 7M screen (In-Stock_Screen.xlsx)
- ID’s, SMILES, Docking scores, % displacement and K_i_ values of tested compounds from the 1.6B screen (1.6B_Screen.xlsx)
- ID’s, SMILES, Docking scores, % displacement and K_i_ values of tested compounds from the 2.6B screen (2.6B_Screen.xlsx)
- Synthetic procedures (Synthetic_procedures.docx), and LC-MS and NMR spectra (Spectra.docx) of described compounds

## AUTHOR INFORMATION

### Authors

**Moira M. Rachman** – Department of Pharmaceutical Chemistry, University of California, San Francisco, 1700 4th St., Byers Hall Suite 508D, San Francisco, CA 94158, USA

**Christos Iliopoulos-Tsoutsouvas –** Center for Drug Discovery and Department of Pharmaceutical Sciences, Northeastern University, Boston, MA 02115, USA

**Michael D. Sacco –** Department of Molecular and Cellular Physiology, Stanford University School of Medicine, Stanford, CA 94305, USA

**Xinyu Xu** – Department of Pharmaceutical Chemistry, University of California, San Francisco, 1700 4th St., Byers Hall Suite 508D, San Francisco, CA 94158, USA

**Cheng-Guo Wu** – Department of Molecular and Cellular Physiology, Stanford University School of Medicine, Stanford, CA 94305, USA

**Emma Santos** – Department of Molecular and Cellular Physiology, Stanford University School of Medicine, Stanford, CA 94305, USA

**Isabella S. Glenn** – Department of Pharmaceutical Chemistry, University of California, San Francisco, 1700 4th St., Byers Hall Suite 508D, San Francisco, CA 94158, USA

**Lu Paris** – Department of Pharmaceutical Chemistry, University of California, San Francisco, 1700 4th St., Byers Hall Suite 508D, San Francisco, CA 94158, USA

**Michelle K. Cahill** – Department of Anesthesiology, Perioperative and Pain Medicine, Stanford University School of Medicine, Stanford CA 94305 USA

**Suthakar Ganapathy** – Center for Drug Discovery and Department of Pharmaceutical Sciences, Northeastern University, Boston, MA 02115, USA

**Tia A. Tummino** – Department of Pharmaceutical Chemistry, University of California, San Francisco, 1700 4th St., Byers Hall Suite 508D, San Francisco, CA 94158, USA

**Yurii S. Moroz** – Enamine Ltd., Kyiv, 02094, Ukraine; National Taras Shevchenko University of Kyiv, 01601 Kyiv, Ukraine; Chemspace LLC, Kyiv, 02094 Ukraine

**Dmytro S. Radchenko** – Enamine Ltd., Kyiv, 02094, Ukraine

**Meri Okorie** – Department of Pharmaceutical Chemistry, University of California, San Francisco, 1700 4th St., Byers Hall Suite 508D, San Francisco, CA 94158, USA; Graduate Program in Pharmaceutical Sciences and Pharmacogenomics, University of California, San Francisco, San Francisco, CA 94158, USA

**Vivianne Tawfik** – Department of Anesthesiology, Perioperative and Pain Medicine, Stanford University School of Medicine, Stanford CA 94305 USA

**John J. Irwin** – Department of Pharmaceutical Chemistry, University of California, San Francisco, 1700 4th St., Byers Hall Suite 508D, San Francisco, CA 94158, USA

## Author Contributions

M.M.R. conducted docking screens with assistance from T.A.T. and M.O. Ligand optimization was performed by M.M.R. and C.I.-T. with input from B.K.S. and A.M. C.I.-T. and S.G. performed binding assays. M.D.S. performed functional assays (BRET) and prepared the CB2–Gi complex, collected cryo-EM data, and modeled the structure. M.K.C. and VT consulted on physiological relevance of the new cannabinoids. X.X. performed functional assays (GloSensor). C.G.W. performed functional assays (BRET) with assistance from E.S. I.S.G. and L.P. tested compounds for colloidal aggregation. Y.S.M. and D.S.R. supervised synthesis of Enamine compounds. J.J.I. developed the docking libraries used here, working with M.M.R. B.K.S. and M.M.R. wrote the manuscript with input from all authors. B.K.S., A.M. and G.S. conceived and supervised the project.

## Acknowledgements

We thank Brian Kobilka for his support of this project.

## Notes

The authors declare the following competing financial interests(s): B.K.S. is co-founder of Epiodyne, BlueDolphin, and Deep Apple Therapeutics, and serves on SAB for Schrodinger LLC, Vilya Therapeutics, Frontier Discovery Ltd, and on the SRB of Genentech. G.S. and J.J.I are co-founders of and consultants for Deep Apple Therapeutics. No other authors declare competing financial interests.

## TOC Graphic

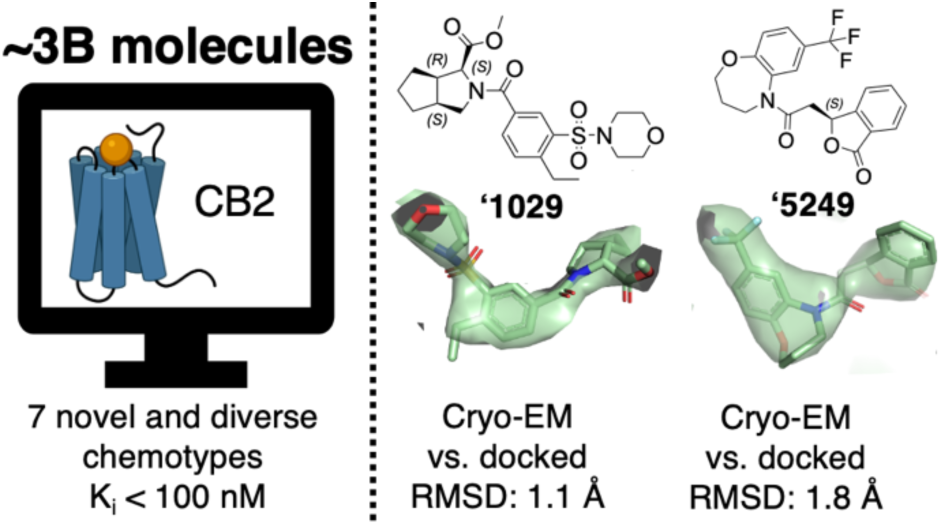

