## Supporting Information for "Library docking for Cannabinoid-2 Receptor ligands"

Contents of SI:

1. Retrieval of known CB2 ligands vs. presumed inactive molecules with the agonist bound docking model.
2. Docking model optimization for more polar molecules.
3. Dynamic light scattering and enzyme inhibition of select compounds from the 7M screen.
4. CB1 and CB2 binding curves of molecules from the 7M screen.
5. CB1 and CB2 binding curves of molecules with $K_i$'s better than 10 μM from the 1.6B screen.
6. CB1 and CB2 binding curves and CB1 3-point data of molecules with $K_i$'s greater than 10 μM from the 1.6B screen.
7. Functional assays ($G_i$ activation) against CB2 of agonists from the 1.6B screen.
8. Functional assays (cAMP accumulation) against CB2 of an inverse agonist from the 1.6B screen.
9. Retrieval of known CB2 ligands vs. presumed inactive molecules with the inverse agonist bound docking model.
10. CB2 binding curves of molecules from the 2.6B screen.
11. Functional assays (cAMP accumulation) against CB2 of agonists and inverse agonists from the 2.6B screen.
12. Functional assays ($G_i$ activation) against CB2 of analogues of select agonists.
13. CB1 and CB2 binding curves of select analogs of an in-stock library hit.
14. Functional assays ($G_i$ activation) against CB2 of stereochemically purified analogs ('2167).
15. Functional assays ($G_i$ activation) against CB2 of stereochemically purified analogs ('5249).
16. CB1R binding curves of compounds <100 nM.
17. Sanity check of the agonist bound model in retrieving neutral over charged molecules.
18. Synthetic procedures.
19. In-Stock_Screen.xls: Excel file listing compounds tested, and their activities, from the 7 million compound "in-stock" docking screen (separate file)
20. 1.6B_screen.xlsx: Excel file listing compounds tested, and. their activities from the 1.6 billion compound docking screen (separate file).
21. 2.6B_Screen.xlsx: Excel file listing compounds tested, and their activities from the 2.6 billion compound docking screen (separate file).

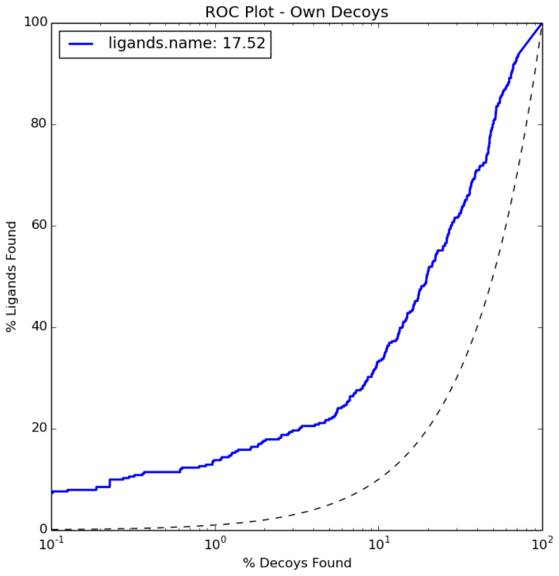


**Figure S1**. Enrichment (retrieved actives versus decoys) with the optimized CB2 agonist bound structure (6PT0). Actives comprised 34 CB2 ligands from the IUPHAR database. Decoys were property-matched but topologically distinct from ligands. Decoys were obtained from the DUDE-Z database.

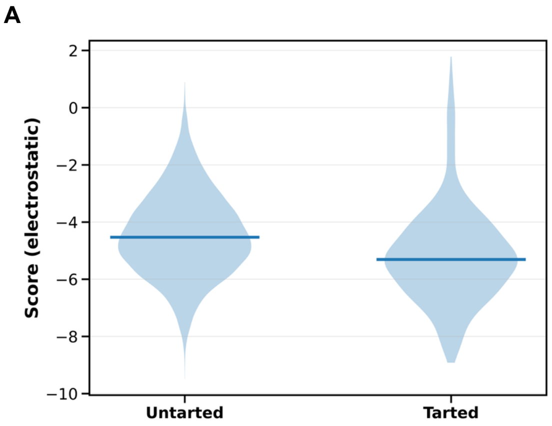


**B**

| | Untarted | Tarted | cLogP | SMILES |
|---|---|---|---|---|
| AM2201 | -6.262029 | -6.393898 | 5.5498 | O=C(C1=CN(CCCCCF)C2=CC=CC=C12)C1=C2C=CC=CC2=CC=C1 |
| AM7499 | -1.5205 | -1.569149 | 5.2347 | CCCCOC(=O)C(C)(C)C1=CC(O)=C2C(=C1)OC(C)(C)C1CCC(CO)CC21 |
| HU-210 | -3.512889 | -3.402883 | 6.9355 | CCCCCCC(C)(C)C1=CC(O)=C2C(=C1)OC(C)(C)C1CC=C(CO)CC21 |
| HU-243 | -1.196183 | -1.189381 | 7.04 | CCCCCCC(C)(C)C1=CC(O)=C2C(=C1)OC(C)(C)C1CCC(CO)CC21 |
| JWH-018 | -6.357453 | -6.238442 | 5.7754 | CCCCCN1C=C(C(=O)C2=C3C=CC=CC3=CC=C2)C2=CC=CC=C21 |

**Figure S2.** More polar molecules were sought by scaling electrostatic potential grids. A model without scaled electrostatics (“Untarted”) showed worse electrostatic scores than a model with scaled electrostatics (“Tarted”) for IUPHAR CB2 ligands **(A)**. For IUPHAR ligands that gave the same pose between these models, less lipophilic ligands (AM2201, AM7499) obtained better electrostatic scores in the “Tarted” model, while more lipophilic ligands (HU-210, HU-243 and JWH-018) showed a worse score **(B)**.

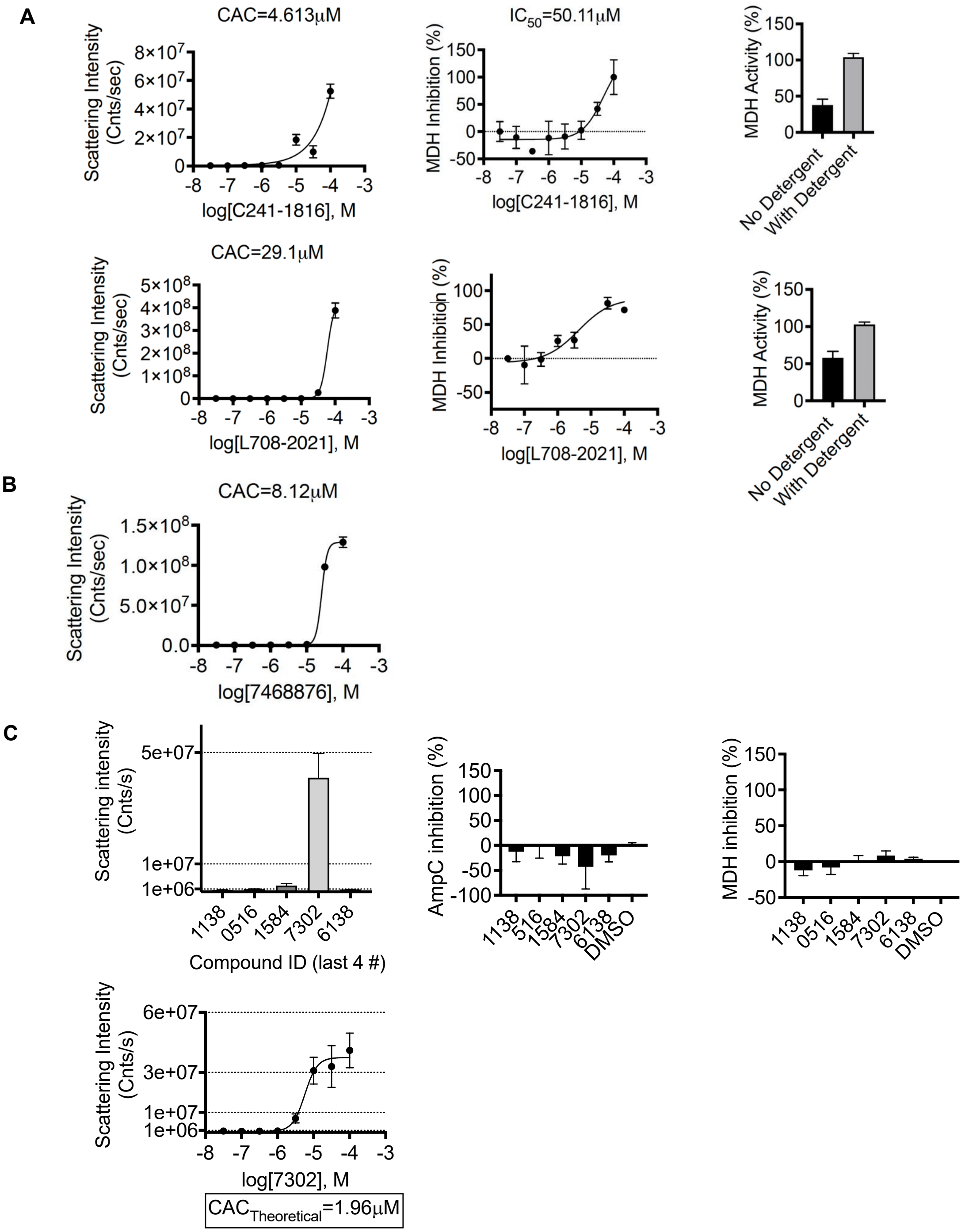


**Figure S3**. Colloidal aggregation testing on eight in-stock screen hits (at least 30% displacement at 10 μM). '1816 and '2021 formed particles and inhibited the counter-enzymes MDH in a detergent dependent manner **(A)**. '8876 formed particles, but did not inhibit counter-enzymes MDH or AmpC **(B)**. '1138, '0516, '1584 and '6138 did not form particles, nor inhibited MDH or AmpC. '7302 formed particles but did not inhibit MDH or AmpC **(C)**.

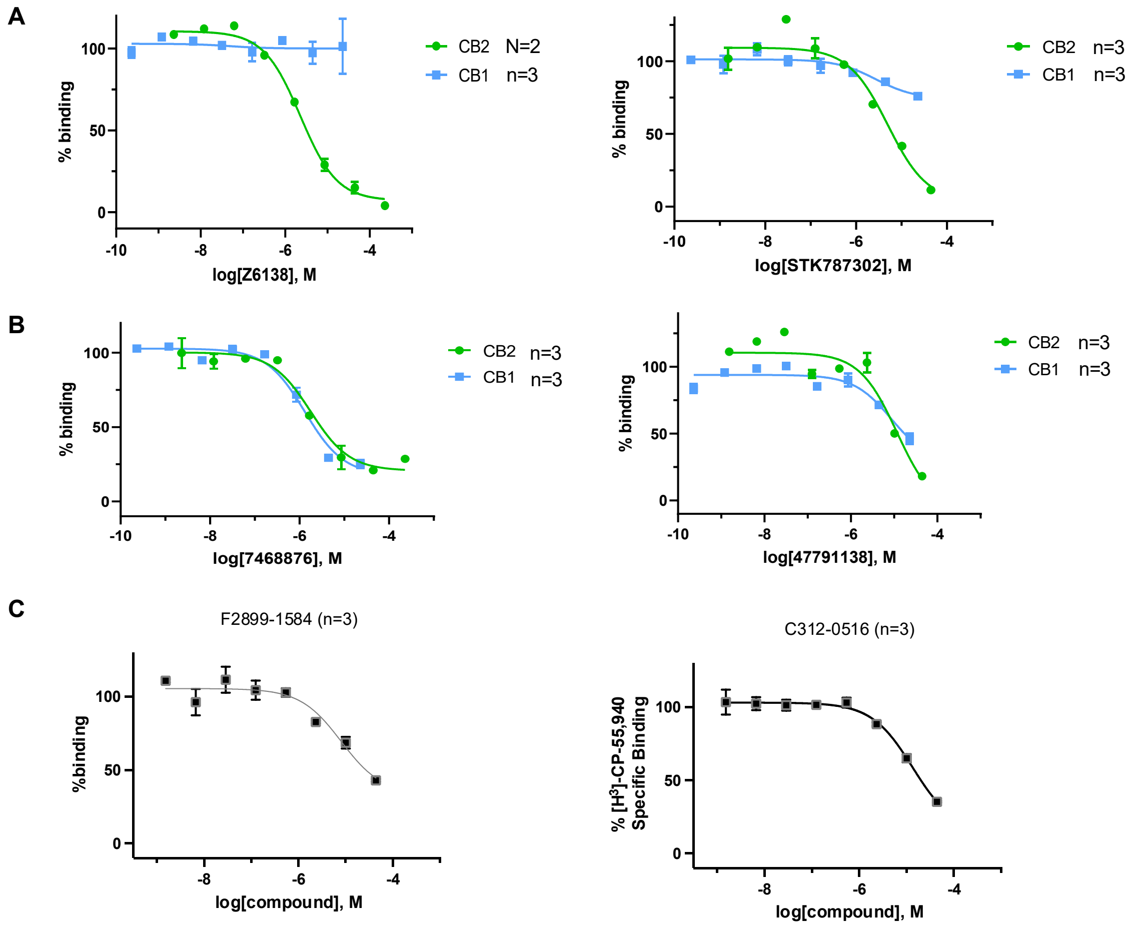


**Figure S4**. Binding curves of in-stock hits that were CB2 selective **(A)**, non-selective **(B)** and not tested for CB1 **(C)**. $K_i$ values were determined from displacement of [$^3$H]-CP-55,940 minus non-specific binding. n = technical repeats, N = independent experiments.

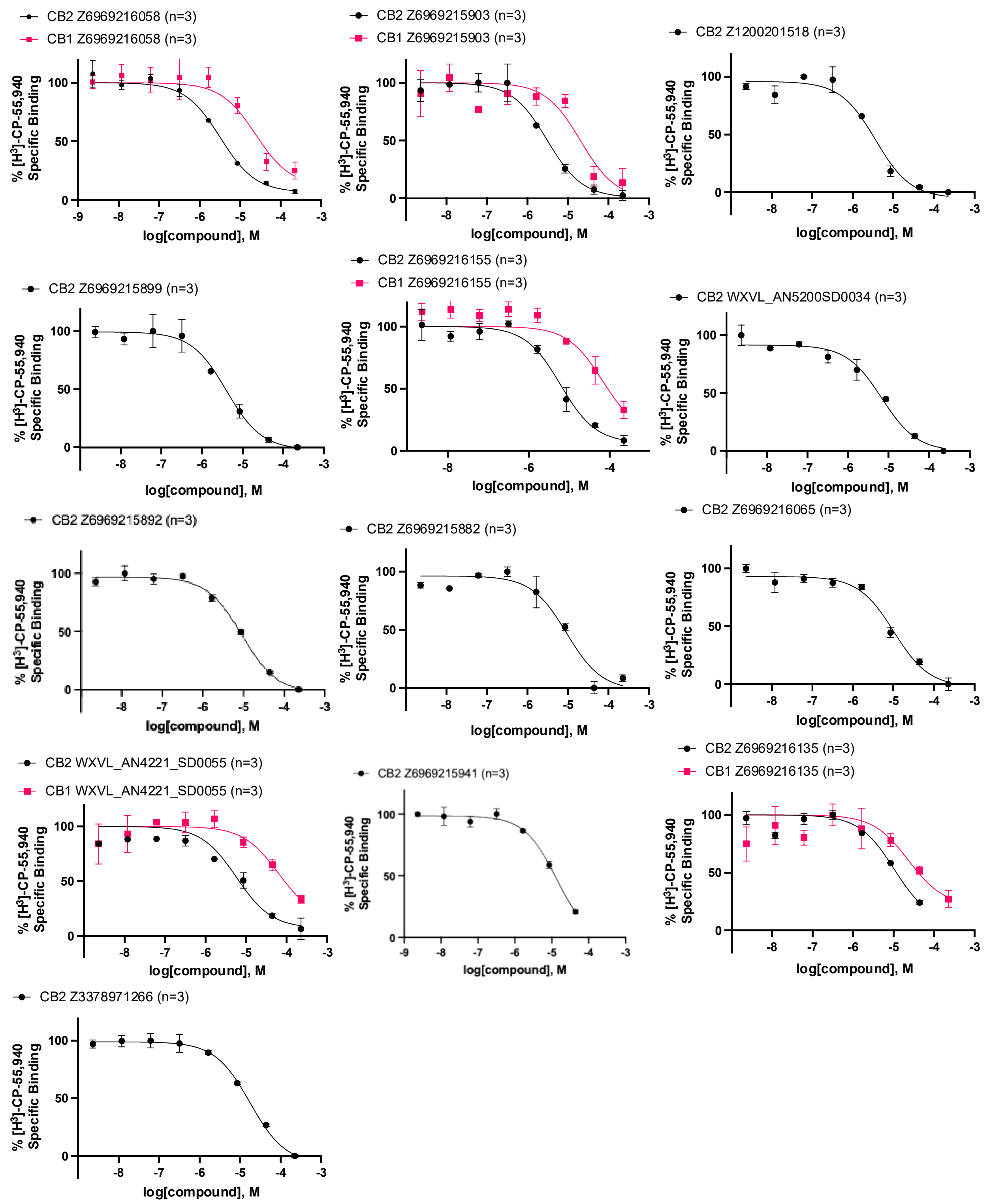


**Figure S5**. Binding curves of CB2 ligands with $K_i$'s <10 μM from a 1.6B library, alongside CB1 curves when tested. $K_i$ values were determined from displacement of [$^3$H]-CP-55,940 minus non-specific binding. n = technical repeats, N = independent experiments.

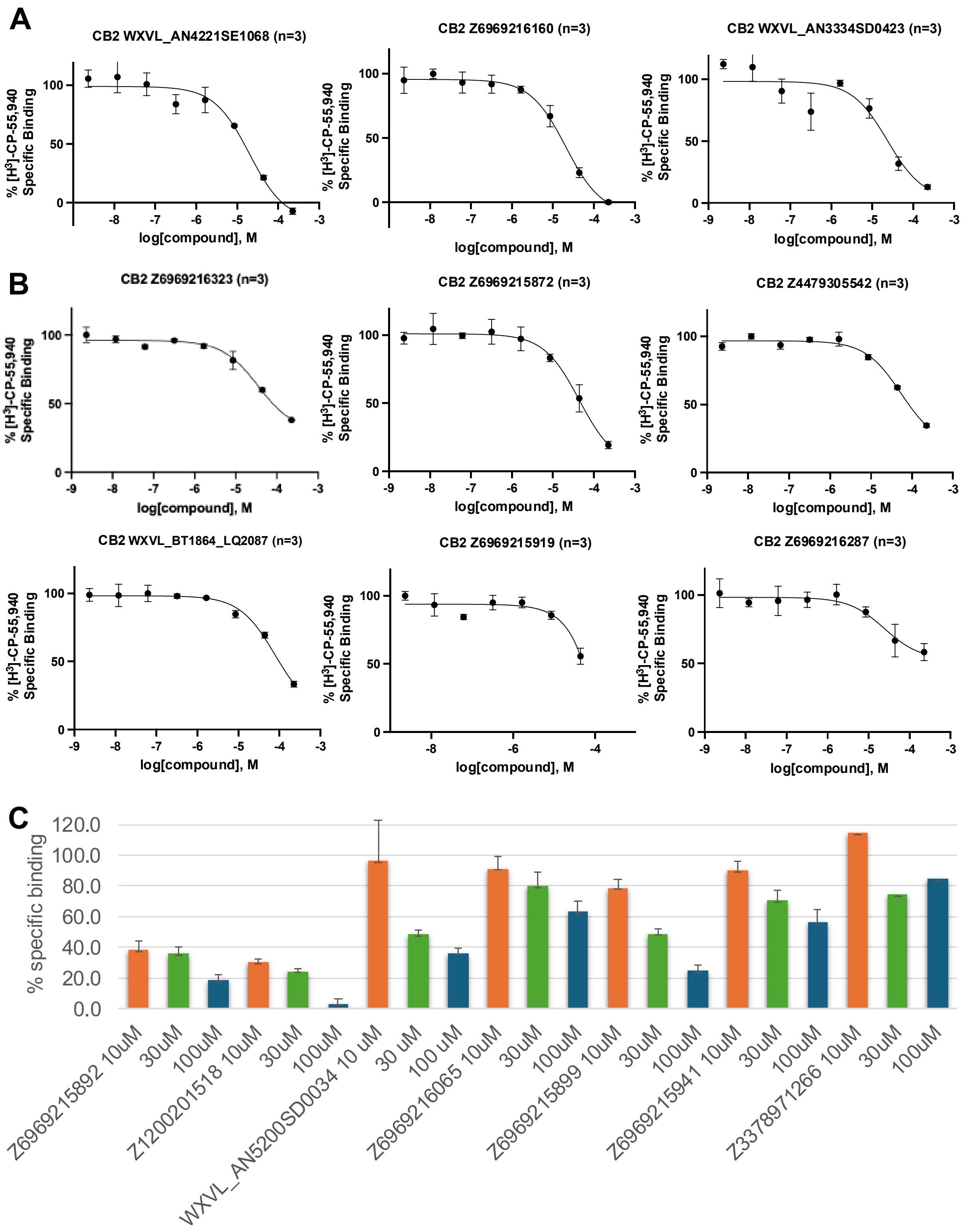


**Figure S6**. Binding curves of CB2 ligands with 10 μM < $K_i$'s < 15 μM **(A)**, $K_i$'s >15 μM **(B)** and CB1 binding at 10 μM, 30 μM and 100 μM of 7 CB2 ligands **(C)** from a 1.6B library. $K_i$ values were determined from displacement of [$^3$H]-CP-55,940 minus non-specific binding. n = technical repeats, N = independent experiments.

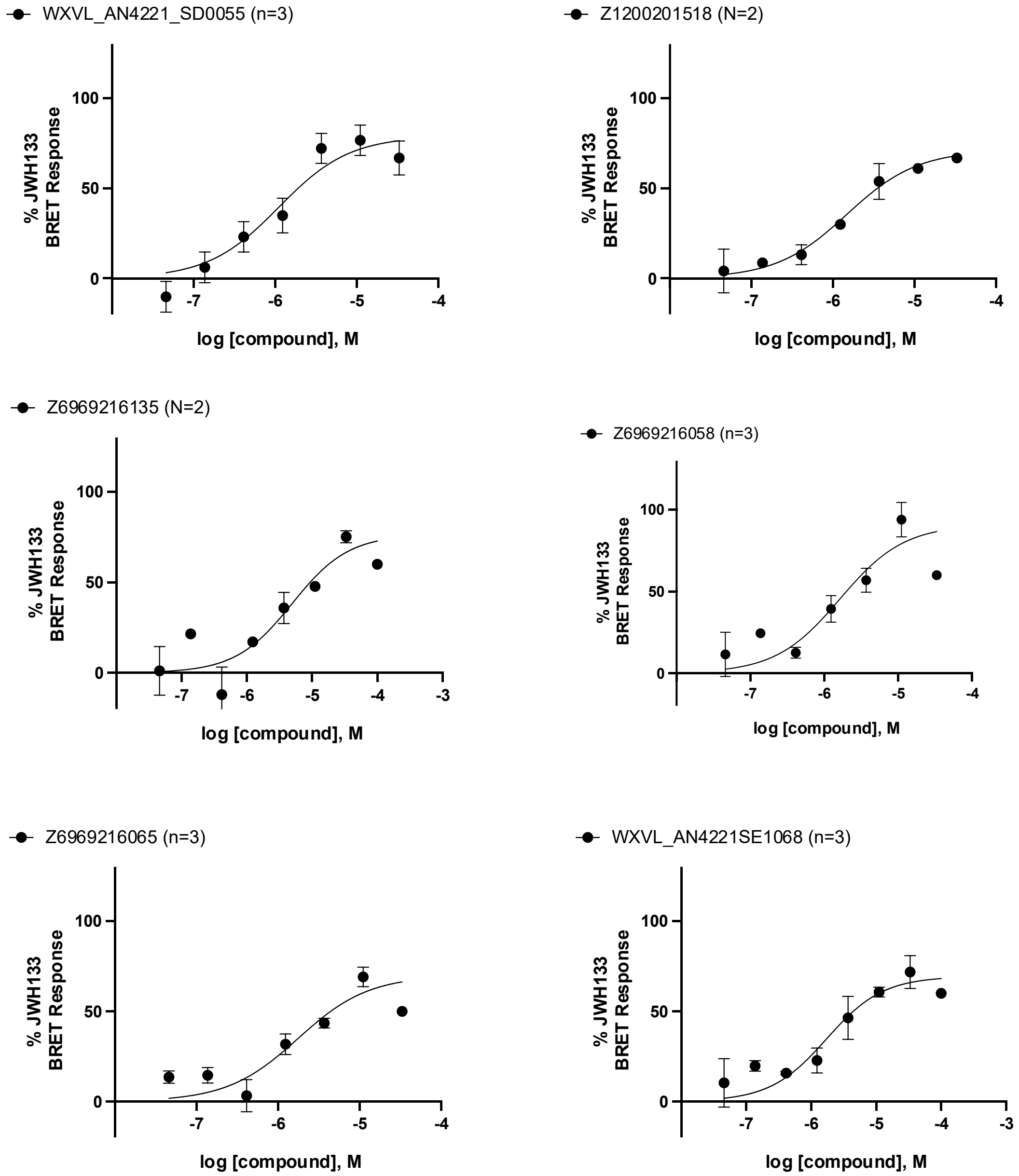


**Figure S7.** Functional activity of six CB2 agonists from a 1.6B library. Agonism was determined from BRET signals monitoring $G_i$ activation normalized to JWH133 signals. n = technical repeats, N = independent experiments.

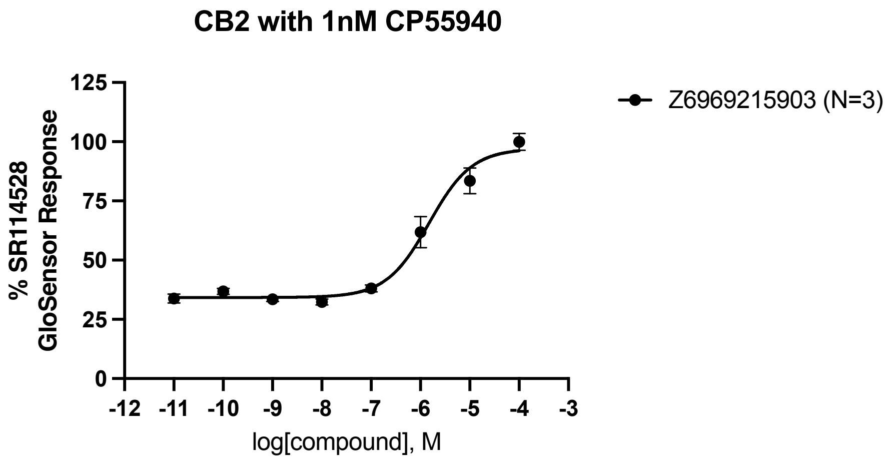


**Figure S8.** Functional activity of the CB2 inverse agonist from a 1.6B library. Inverse agonism was determined by monitoring cAMP accumulation with GloSensor in the presence of Forskolin and CP-55,940 normalized to SR114528. n = technical repeats, N = independent experiments.

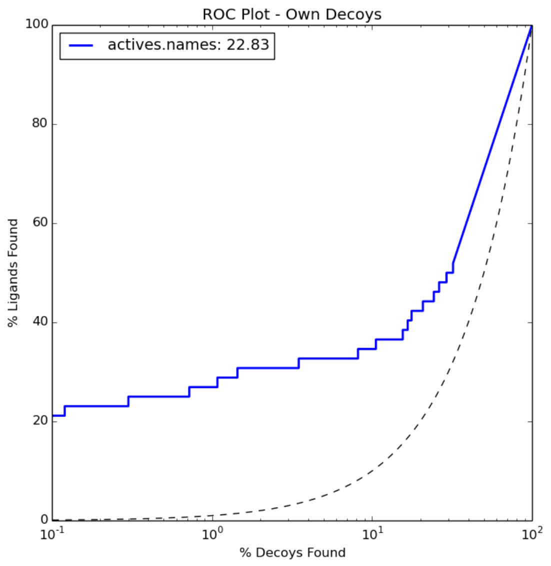


**Figure S9**. Enrichment (retrieved actives versus decoys) with the optimized CB2 inverse agonist bound structure (5ZTY). Actives comprised 34 CB2 ligands from the IUPHAR database. Decoys were property-matched but topologically distinct from ligands. Decoys were obtained from the DUDE-Z database.

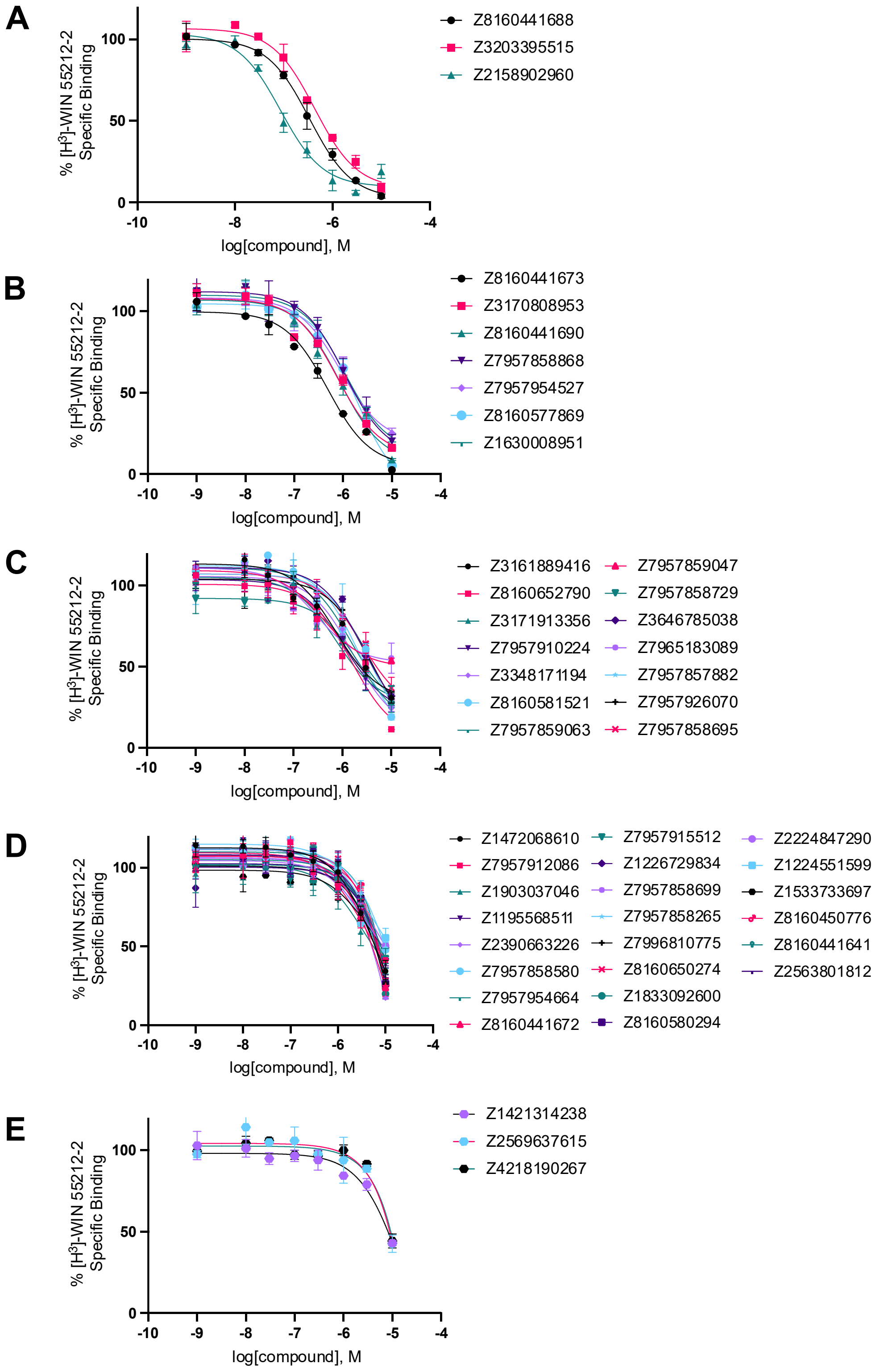


**Figure S10.** CB2 $K_i$ curves 30 nM - 300 nM **(A)**, 300 nM - 1 µM **(B)**, 1 µM – 3 µM **(C)**, 3 µM – 10 µM **(D)**, and 10 µM – 15 µM **(E)** of molecules from the 2.6B screen that showed >30% displacement. All points are from technical duplicates or n=2. $K_i$ values were determined from displacement of [$^3$H]- WIN 55212-2 minus non-specific binding.

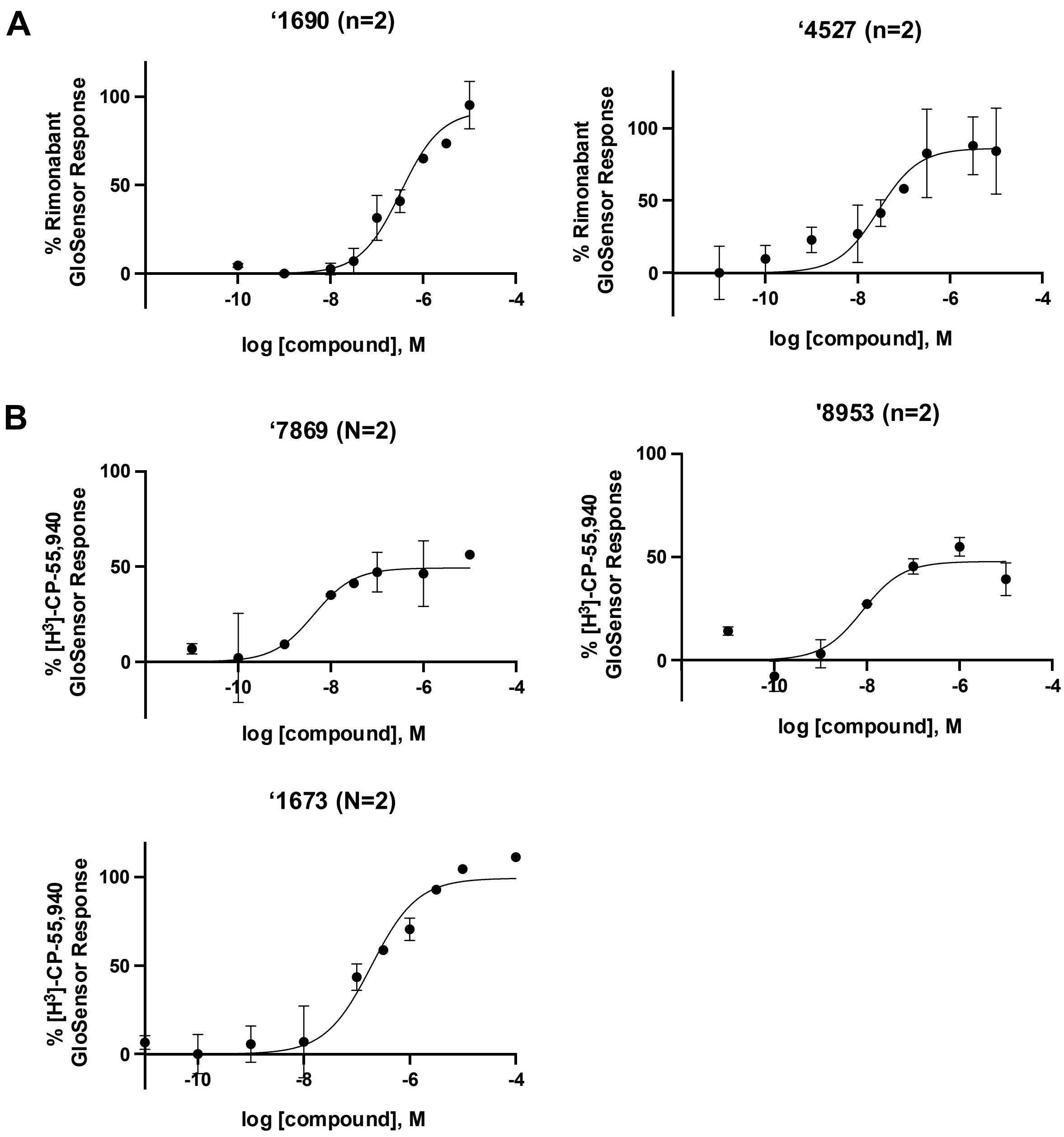


**Figure S11.** GloSensor curves of 2 inverse agonists **(A)** and 3 agonists **(B)** from the 2.6B screen. Inverse agonism was determined by monitoring cAMP accumulation with GloSensor in the presence of Forskolin and CP-55,940 normalized to Rimonabant. Agonism was determined by monitoring cAMP accumulation with GloSensor in the presence of Forskolin normalized to CP-55,940. n = technical repeats, N = independent experiments.

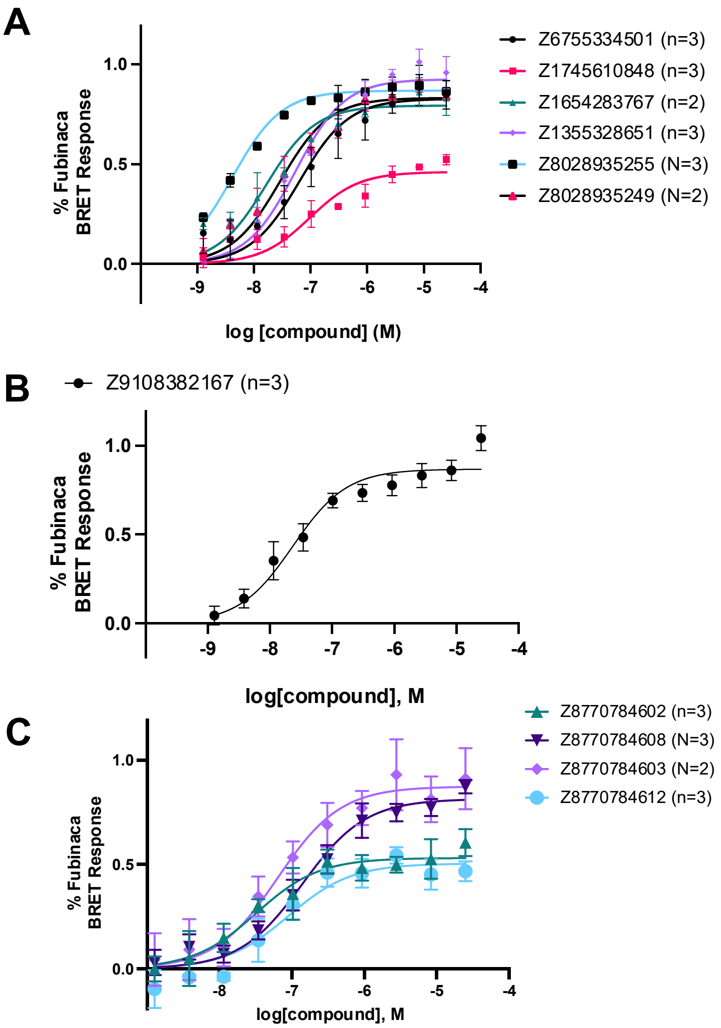


**Figure S12.** Activity of representative analogs for 3 compounds, ‘6138 from the 7M screen **(A)**, ‘1068 from the 1.6B screen **(B)** and ‘7302 from the 7M screen **(C)**. Agonism was determined from BRET signals monitoring $G_i$ activation normalized to Fubinaca signals. n = technical repeats, N = independent experiments.

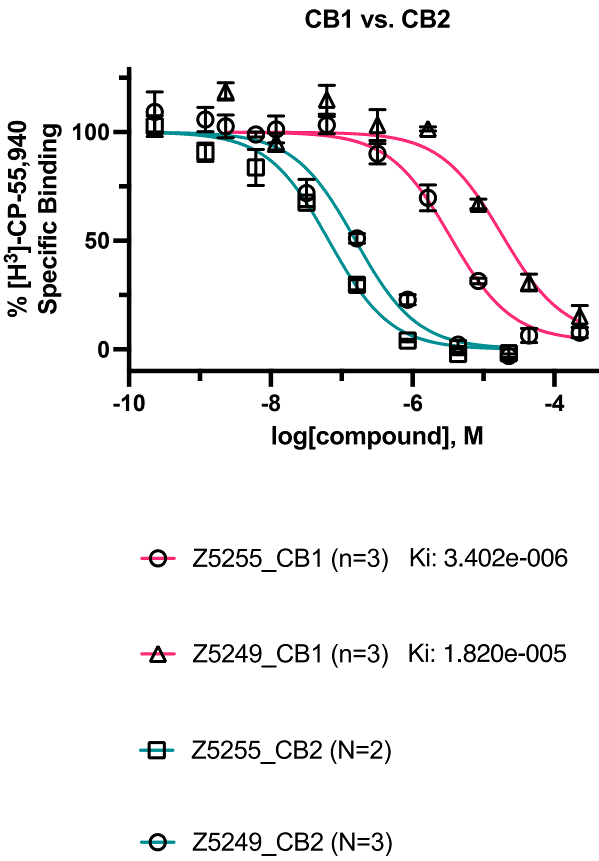


**Figure S13.** Analogs of '6138 from the 7M library screen, '5249 has 115-fold selectivity, while '5255 has 51-fold selectivity over CB1. $K_i$ values were determined from displacement of [$^3$H]-CP-55,940 minus non-specific binding. n = technical repeats, N = independent experiments.

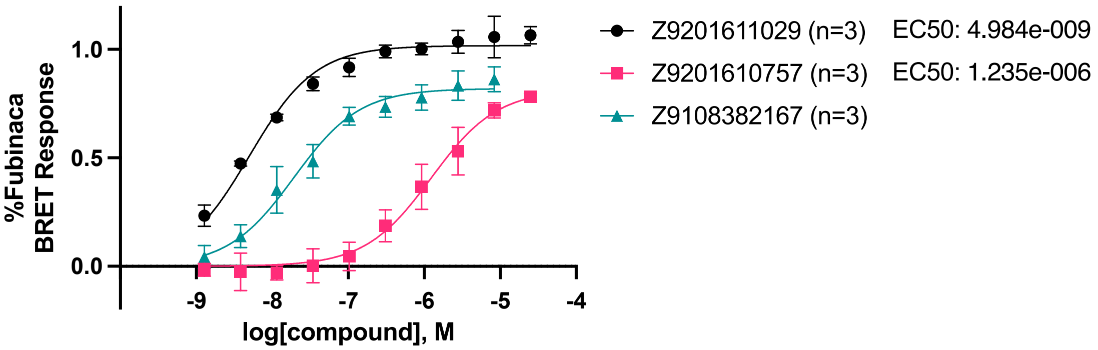


**Figure S14.** Functional activity of stereochemically purified '2167 cis isomers. Agonism was determined from BRET signals monitoring $G_i$ activation normalized to Fubinaca signals. Purification was performed at Enamine. n = technical repeats, N = independent experiments.

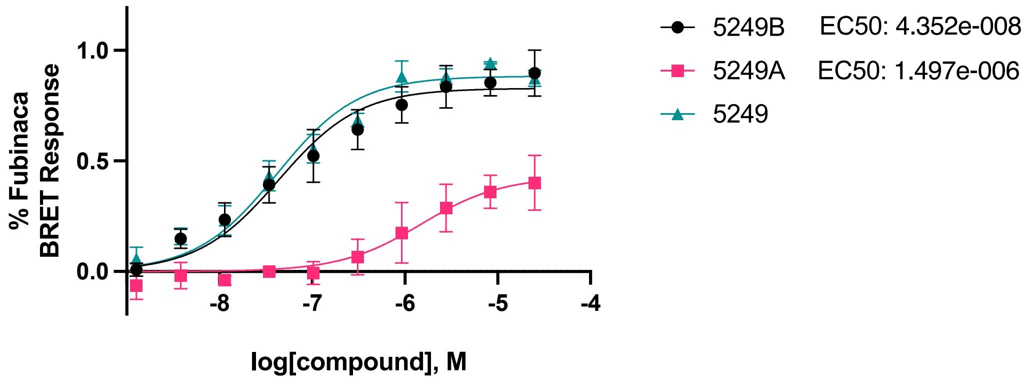


**Figure S15.** Purified '5249 showed the better isomer being in the same activity range as the racemic mixture. Agonism was determined from BRET signals monitoring $G_i$ activation normalized to Fubinaca signals. Purification was performed at Northeastern University. Data points are from technical triplicates.

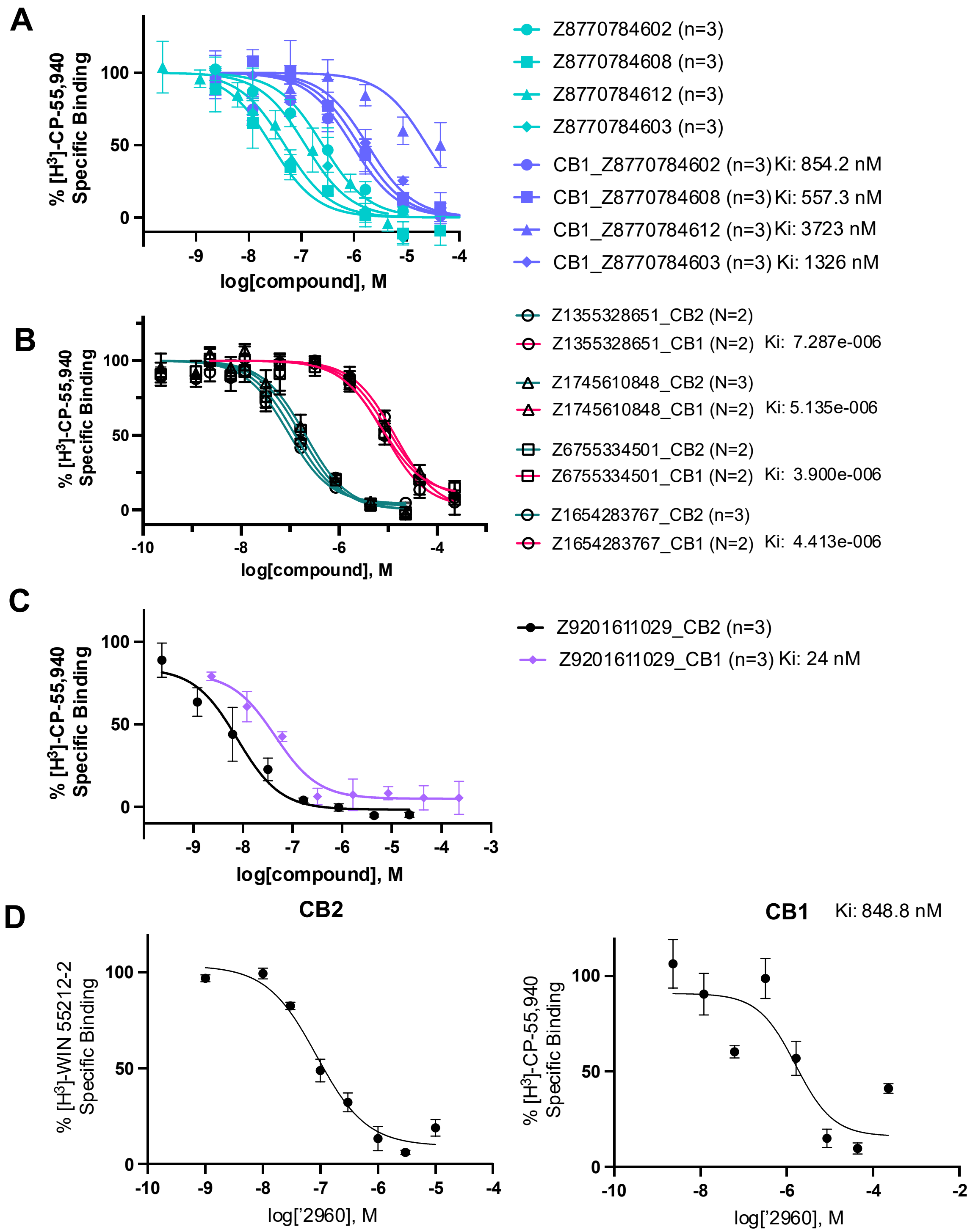


**Figure S16.** Selectivity over CB1R of discovered compounds <100 nM. $K_i$ values were determined from displacement of [$^3$H]-CP-55,940 or [$^3$H]- WIN 55212-2 minus non-specific binding. n = technical repeats, N = independent experiments.

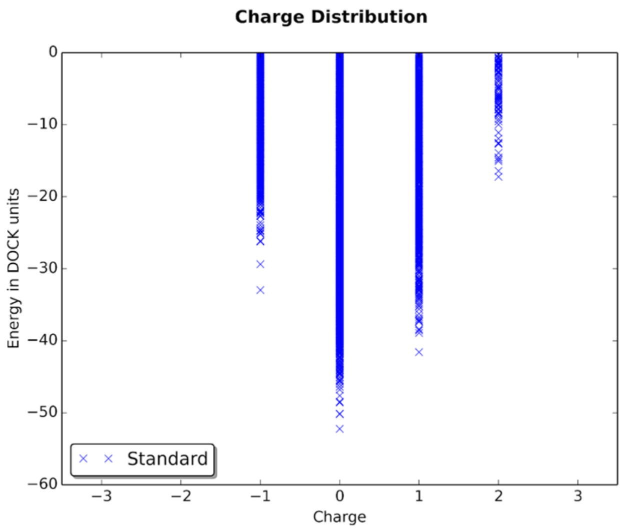


**Figure S17.** Sanity check of the agonist bound model in retrieving neutral over charged molecules.

**Synthetic procedures.**

**General considerations**. All chemicals for the synthesis of the studied compounds were provided by Enamine Ltd. (www.enamine.net). All solvents were treated according to standard methods. $^1$H NMR spectra were recorded at 400, 500, or 600 MHz (Varian or Bruker spectrometers). $^1$H chemical shifts are calibrated using residual nondeuterated solvent DMSO, δ = 2.50 ppm. Coupling constants are given in Hz. LC/MS analysis was performed utilizing an Agilent 1200 Series LC/MSD system with DAD/ELSD (column Zorbax SB-C18 1.8 µm 4.6x15 mm; solvent A (water, 0.1% formic acid) and solvent B (acetonitrile, 0.1% formic acid); gradient 0% – 100% solvent B, run time, 1.8 min; flow rate, 3 mL/min) and Agilent LC/MSD SL (G6130A) or SL (G6140A) mass spectrometer (APCI mode). All the LC/MS data were obtained using positive/negative mode switching.

**Method 1.** The synthesis was performed according to a previously published procedure (dx.doi.org/10.1021/acscombsci.6b00103).

**Method 2.** A carboxylic acid (100 mg) was dissolved in 0.5 mL of 10% Hydroxybenzotriazole (HOBt) in DMF, followed by the addition of an amine (1 mol equiv to the carboxylic acid) and 1-ethyl-3-(3-dimethylaminopropyl)carbodiimide (EDC) (1.2 mol equiv to the carboxylic acid). The resulting mixture was shaken for 24 h at room temperature. Then, $CHCl_3$ (2 mL) was added, and the organic phase was washed with water, dried over sodium sulfate, and evaporated under reduced pressure. The crude was dissolved in 0.5 mL of DMSO and further purified by preparative HPLC.

**Method 3.** A carboxylic acid (100 mg) was dissolved in 0.5 mL of DMF, followed by the addition of an alcohol (1 mol equiv to the carboxylic acid) and 1-ethyl-3-(3-dimethylaminopropyl)carbodiimide (EDC) (1.2 mol equiv to the carboxylic acid) and 4-*N,N*-dimethylaminopyridine (DMAP) (0.05 mol equiv to the carboxylic acid). The resulting mixture was shaken for 24 h at room temperature. Then, $CHCl_3$ (2 mL) was added, and the organic phase was washed with water, dried over sodium sulfate, and evaporated under reduced pressure. The crude was dissolved in 0.5 mL of DMSO and further purified by preparative HPLC.

**Method 4.** The synthesis was performed according to the previously published procedure (dx.doi.org/10.1016/j.tetlet.2010.06.139).

**Method 5.** The synthesis was performed according to the previously published procedure (dx.doi.org/10.1007/s11030-021-10218-2).

**Method 6.** The synthesis was performed according to the previously published procedure (dx.doi.org/10.1021/co500025f).

**Method 7.** An amine (50 mg) was dissolved in 0.5 mL of glacial acetic acid in a 4 mL vial. Then sodium acetate (1.1 mol equiv to the amine) and a sulfonyl chloride (1 mol equiv to the amine) were added. The vial was sealed and kept in an ultrasonic bath for 24 h. Then, the crude sample was dissolved in 0.5 mL of DMSO and purified by preparative HPLC.

**Method 8.** The synthesis was performed according to the previously published procedure (dx.doi.org/10.1055/s-1984-30909 or dx.doi.org/10.1002/jhet.5570240324).

**Method 9.** An amine (100 mg), DIPEA (1.2 mol equivalent to the amine), and DMSO (0.5 mL) were placed into a 4 mL capped glass vial and stirred for 30 min. After the addition of an alkyl halide (1.2 mol equiv to the amine), the vial was stirred for 12 h at room temperature. Then the vial was placed into a thermostat (set to 100°C) for 9 h. After cooling down, the mixture was filtered; the solvent and volatile components were evaporated under reduced pressure to give the crude product. The product was further purified by HPLC.

**Spectral description of the synthesized compounds.**

**5-bromo-8-[5-(2-chloro-2-fluoro-1-methylcyclopropyl)-1,2,4-oxadiazol-3-yl]isoquinoline – Z8160577869 (Method 1).**
Yield: 20%; purity, >95% (assessed by LC/MS).
LC/MS (APSI) m/z [M+H] calculated for C15H11BrClFN3O: 382.0; found: 382.0.

**3-(4-bromo-2-methoxyphenyl)-5-(2-cyclopropoxyphenyl)-1,2,4-oxadiazole – Z2158902960 (Method 1).**
Yield: 29%; purity, >95% (assessed by LC/MS).
LC/MS (APSI) m/z [M+H] calculated for C18H16BrN2O3: 389.0; found: 389.0.

**2-(cyclobutylmethyl)-1-[5-(2-methanesulfonylpropan-2-yl)-1,3,4-oxadiazole-2-carbonyl]piperidine – Z3203395515 (Method 2).**
Yield: 16%; purity, >95% (assessed by LC/MS).
LC/MS (APSI) m/z [M+H] calculated for C17H28N3O4S: 370.2; found: 370.4.

**1-[2,5-dimethyl-1-(1,3-thiazol-2-yl)-1H-pyrrol-3-yl]-2-[(2R)-2-ethynyl-2-methylpyrrolidin-1-yl]ethan-1-one trifluoroacetate – Z8160441688 (Method 9).**
Yield: 28%; purity, >95% (assessed by LC/MS).
LC/MS (APSI) m/z [M+H] calculated for C18H22N3OS: 328.2; found: 328.0.

**(2E)-1-(7-fluoro-6-methoxy-1,2,3,4-tetrahydroquinolin-1-yl)-4-(4-methylphenyl)but-2-ene-1,4-dione – Z8160441641 (Method 2).**
Yield: 40%; purity, >95% (assessed by LC/MS).
LC/MS (APSI) m/z [M+H] calculated for C21H21FNO3: 354.2; found: 354.0.

**methyl 2-[5-bromo-1-(2,2-difluoro-4-methylpentanoyl)-2,3-dihydro-1H-indol-3-yl]acetate – Z7957910224 (Method 2).**
Yield: 24%; purity, >95% (assessed by LC/MS).
LC/MS (APSI) m/z [M+H] calculated for C17H21BrF2NO3: 404.1; found: 404.2.

**(2-bromo-2-methylcyclopropyl)methyl 6-(4-fluorophenyl)pyridine-2-carboxylate – Z7957954664 (Method 3).**
Yield: 33%; purity, >95% (assessed by LC/MS).
LC/MS (APSI) m/z [M+H] calculated for C17H16BrFNO2: 366.0; found: 366.0.

**2-[4-(cyclopropylmethyl)-2-methyl-4H-thieno[3,2-b]pyrrol-5-yl]-5-[(2S)-oxolan-2-yl]-1,3,4-oxadiazole – Z3348171194 (Method 4).**

Yield: 29%; purity, >95% (assessed by LC/MS).
LC/MS (APSI) m/z [M+H] calculated for C17H20N3O2S: 330.1; found: 330.1.

**5-chloro-1-(6-fluoro-3,4-dihydro-2H-1-benzopyran-2-carbonyl)-3,3-dimethyl-2,3-dihydro-1H-indole – Z7957915512 (Method 2).**
Yield: 58%; purity, >95% (assessed by LC/MS).
LC/MS (APSI) m/z [M+H] calculated for C20H20ClFNO2: 360.1; found: 360.0.

**(3-cyanothiophen-2-yl)methyl 1-phenyl-5-(propan-2-yl)-1H-pyrazole-3-carboxylate – Z7957858868 (Method 3).**
Yield: 29%; purity, >95% (assessed by LC/MS).
LC/MS (APSI) m/z [M+H] calculated for C19H18N3O2S: 352.1; found: 352.2.

**6-bromo-8-methyl-2-{4H,5H,6H,7H-thieno[3,2-c]pyridine-5-carbonyl}imidazo[1,2-a]pyridine – Z8160441672 (Method 2).**
Yield: 24%; purity, >95% (assessed by LC/MS).
LC/MS (APSI) m/z [M+H] calculated for C16H15BrN3OS: 376.0; found: 375.9.

**1-{bicyclo[4.2.0]octa-3,7-diene-7-carbonyl}-5-chloro-3,3-dimethyl-2,3-dihydro-1H-indole – Z8160580294 (Method 2).**
Yield: 14%; purity, >95% (assessed by LC/MS).
LC/MS (APSI) m/z [M+H] calculated for C19H21ClNO: 314.1; found: 314.0.

**4-(7-fluoro-4-methyl-1-benzofuran-2-carbonyl)-2-methyl-6-(thiophen-3-yl)morpholine – Z8160650274 (Method 2).**
Yield: 62%; purity, >95% (assessed by LC/MS).
LC/MS (APSI) m/z [M+H] calculated for C19H19FNO3S: 360.1; found: 360.0.

**{4-methyl-2-oxabicyclo[2.2.1]heptan-1-yl}methyl 3-bromo-5-cyclopropoxybenzoate – Z7957954527 (Method 3).**
Yield: 18%; purity, >95% (assessed by LC/MS).
LC/MS (APSI) m/z [M+H] calculated for C18H22BrO4: 381.1; found: 381.0.

**N-[5-(cyclopropylmethyl)-1,2-oxazol-3-yl]-5-fluoro-2-(propan-2-yloxy)benzamide – Z8160441690 (Method 2).**
Yield: 24%; purity, >95% (assessed by LC/MS).
LC/MS (APSI) m/z [M+H] calculated for C17H20FN2O3: 319.1; found: 319.2.

**(3-fluoropyridin-4-yl)methyl 3-methoxy-5-phenylthiophene-2-carboxylate – Z7957858699 (Method 3).**
Yield: 27%; purity, >95% (assessed by LC/MS).
LC/MS (APSI) m/z [M+H] calculated for C18H15FNO3S: 344.1; found: 344.0.

**(3,5-dimethyl-1,2-oxazol-4-yl)methyl 1-phenyl-5-(propan-2-yl)-1H-pyrazole-3-carboxylate – Z1472068610 (Method 3).**
Yield: 42%; purity, >95% (assessed by LC/MS).
LC/MS (APSI) m/z [M+H] calculated for C19H22N3O3: 340.2; found: 340.2.

**(1-ethyl-1H-imidazol-5-yl)methyl 3-methoxy-5-phenylthiophene-2-carboxylate triflate – Z7957858695 (Method 3).**
Yield: 29%; purity, >95% (assessed by LC/MS).
LC/MS (APSI) m/z [M+H] calculated for C18H19N2O3S: 343.1; found: 343.2.

**(2-bromo-2-methylcyclopropyl)methyl 3-phenyl-5H,6H,8H-imidazo[4,3-c][1,4]oxazine-1-carboxylate – Z3170808953 (Method 3).**
Yield: 22%; purity, >95% (assessed by LC/MS).
LC/MS (APSI) m/z [M+H] calculated for C18H20BrN2O3: 391.1; found: 391.0.

**(1-methylcyclobutyl)methyl 3-phenyl-5H,6H,8H-imidazo[4,3-c][1,4]oxazine-1-carboxylate – Z3161889416 (Method 3).**
Yield: 34%; purity, >95% (assessed by LC/MS).
LC/MS (APSI) m/z [M+H] calculated for C19H21N2O3: 327.2; found: 327.2.

**1-(pyridin-3-yl)ethyl 1-phenyl-1H,4H,5H,6H-cyclopenta[c]pyrazole-3-carboxylate – Z1195568511 (Method 3).**
Yield: 25%; purity, >95% (assessed by LC/MS).
LC/MS (APSI) m/z [M+H] calculated for C20H20N3O2: 334.2; found: 334.2.

**2-[4-bromo-2-(propan-2-yloxy)phenyl]-5-(2-methyloxolan-2-yl)-1,3,4-oxadiazole – Z2390663226 (Method 4).**
Yield: 28%; purity, >95% (assessed by LC/MS).
LC/MS (APSI) m/z [M+H] calculated for C16H20BrN2O3: 369.1; found: 369.0.

**1-(prop-2-en-1-yl)-3-[5-(thiophen-3-yl)-1,3,4-oxadiazol-2-yl]-1H-indole – Z8160441673 (Method 4).**
Yield: 30%; purity, >95% (assessed by LC/MS).
LC/MS (APSI) m/z [M+H] calculated for C17H14N3OS: 308.1; found: 308.2.

**(1-cyanocyclopentyl)methyl 7-methoxy-2-methyl-2,3-dihydro-1-benzofuran-5-carboxylate – Z1630008951 (Method 3).**
Yield: 33%; purity, >95% (assessed by LC/MS).
LC/MS (APSI) m/z [M+H] calculated for $C_{18}H_{22}NO_4$: 316.2; found: 316.2.

**(5-methylfuran-3-yl)methyl 5-ethyl-1-(2-fluorophenyl)-1H-1,2,4-triazole-3-carboxylate – Z2224847290 (Method 3).**
Yield: 23%; purity, >95% (assessed by LC/MS).
LC/MS (APSI) m/z [M+H] calculated for $C_{17}H_{17}FN_3O_3$: 330.1; found: 330.2.

**(3-chlorothiophen-2-yl)methyl 5-ethyl-1-phenyl-1H-1,2,4-triazole-3-carboxylate – Z1903037046 (Method 3).**
Yield: 46%; purity, >95% (assessed by LC/MS).
LC/MS (APSI) m/z [M+H] calculated for $C_{16}H_{15}ClN_3O_2S$: 348.1; found: 348.0.

**1-(3,3-dimethylcyclopentyl) 3-methyl 4,5,6,7-tetrahydro-2-benzothiophene-1,3-dicarboxylate – Z7965183089 (Method 3).**
Yield: 24%; purity, >95% (assessed by LC/MS).
LC/MS (APSI) m/z [M-H] calculated for $C_{18}H_{25}O_4S$: 335.1; found: 335.2.

**5-(1-tert-butyl-1H-1,2,4-triazol-3-yl)-3-cyclopropyl-1-[(3-methylcyclobutyl)methyl]-1H-1,2,4-triazole – Z8160581521 (Method 5).**
Yield: 15%; purity, >95% (assessed by LC/MS).
LC/MS (APSI) m/z [M+H] calculated for $C_{17}H_{27}N_6$: 315.2; found: 315.2.

**5-(2-benzyl-1,3-thiazol-5-yl)-3-ethyl-1-[(thiolan-3-yl)methyl]-1H-1,2,4-triazole – Z8160652790 (Method 5).**
**Yield: 16%; purity, >95% (assessed by LC/MS).**
LC/MS (APSI) m/z [M+H] calculated for $C_{19}H_{23}N_4S_2$: 371.1; found: 371.0.

**1-(3-fluorophenyl)-5-(propan-2-yl)-3-[2-(propan-2-yl)-1,3-oxazolidine-3-carbonyl]-1H-1,2,4-triazole – Z4218190267 (Method 2).**
Yield: 26%; purity, >95% (assessed by LC/MS).
LC/MS (APSI) m/z [M+H] calculated for $C_{18}H_{24}FN_4O_2$: 347.2; found: 347.0.

**2-ethyl-6-methyl-4-{[5-(2-methylphenyl)-1,2,4-oxadiazol-3-yl]methyl}morpholine triflate – Z1421314238 (Method 3).**

Yield: 15%; purity, >95% (assessed by LC/MS).
LC/MS (APSI) m/z [M+H] calculated for $C_{17}H_{24}N_3O_2$: 302.2; found: 302.0.

**8-[(5-{7,7-difluoro-2-oxabicyclo[4.1.0]heptan-1-yl}-1,2,4-oxadiazol-3-yl)methyl]quinoline – Z2569637615 (Method 1).**
Yield: 17%; purity, >95% (assessed by LC/MS).
LC/MS (APSI) m/z [M+H] calculated for $C_{18}H_{16}F_2N_3O_2$: 344.1; found: 344.0.

**2-[5-(oxan-2-yl)-1,3,4-oxadiazol-2-yl]-1-(2,2,2-trifluoroethyl)-1H-indole – Z2563801812 (Method 4).**
Yield: 16%; purity, >95% (assessed by LC/MS).
LC/MS (APSI) m/z [M+H] calculated for $C_{17}H_{17}F_3N_3O_2$: 352.1; found: 352.0.

**methyl 5-bromo-1-(3-ethynyl-3-fluoroazetidine-1-carbonyl)-2,3-dihydro-1H-indole-3-carboxylate – Z8160450776 (Method 6).**
Yield: 18%; purity, >95% (assessed by LC/MS).
LC/MS (APSI) m/z [M+H] calculated for $C_{16}H_{15}BrFN_2O_3$: 381.0; found: 381.0.

**(2-chloro-1,3-thiazol-5-yl)methyl 1-(2-fluorophenyl)-1H,4H,5H,6H-cyclopenta[c]pyrazole-3-carboxylate – Z1533733697 (Method 3).**
Yield: 25%; purity, >95% (assessed by LC/MS).
LC/MS (APSI) m/z [M+H] calculated for $C_{17}H_{14}ClFN_3O_2S$: 378.1; found: 377.9.

**(cyclohex-3-en-1-yl)methyl 5-ethyl-1-phenyl-1H-1,2,4-triazole-3-carboxylate – Z1224551599 (Method 3).**
Yield: 27%; purity, >95% (assessed by LC/MS).
LC/MS (APSI) m/z [M+H] calculated for $C_{18}H_{22}N_3O_2$: 312.2; found: 312.2.

**8-chloro-4-(3-cyclopentyl-1,2-oxazole-5-carbonyl)-2-methyl-3,4-dihydro-2H-1,4-benzoxazine – Z1833092600 (Method 2).**
Yield: 50%; purity, >95% (assessed by LC/MS).
LC/MS (APSI) m/z [M+H] calculated for $C_{18}H_{20}ClN_2O_3$: 347.1; found: 347.0.

**3-{[(4-fluorophenyl)sulfanyl]methyl}-5-(3-methyl-1-phenyl-1H-pyrazol-5-yl)-1,2,4-oxadiazole – Z7957858265 (Method 1).**
Yield: 14%; purity, >95% (assessed by LC/MS).
LC/MS (APSI) m/z [M+H] calculated for $C_{19}H_{16}FN_4OS$: 367.1; found: 367.0.

**(1S,2R)-2-[3-(5-bromoquinolin-8-yl)-1,2,4-oxadiazol-5-yl]cyclohexan-1-ol – Z7996810775 (Method 1).**
Yield: 18%; purity, >95% (assessed by LC/MS).
LC/MS (APSI) m/z [M+H] calculated for $C_{17}H_{17}BrN_3O_2$: 376.0; found: 375.9.

**(2-chloro-4-fluorophenyl)methyl 5-ethyl-1-phenyl-1H-1,2,4-triazole-3-carboxylate – Z1226729834 (Method 3).**
Yield: 54%; purity, >95% (assessed by LC/MS).
LC/MS (APSI) m/z [M+H] calculated for $C_{18}H_{16}ClFN_3O_2$: 360.1; found: 360.0.

**(1-methylcyclopent-3-en-1-yl)methyl 5-ethyl-1-phenyl-1H-1,2,4-triazole-3-carboxylate – Z7957858580 (Method 3).**
Yield: 23%; purity, >95% (assessed by LC/MS).
LC/MS (APSI) m/z [M+H] calculated for $C_{18}H_{22}N_3O_2$: 312.2; found: 312.2.

**3-bromo-5-{3-cyclopentyl-5-[(oxan-2-yl)methyl]-1H-1,2,4-triazol-1-yl}pyridine – Z7957912086 (Method 5).**
Yield: 21%; purity, >95% (assessed by LC/MS).
LC/MS (APSI) m/z [M+H] calculated for $C_{18}H_{23}BrN_4O$: 393.1; found: 393.2.

**(1-methylcyclopent-2-en-1-yl)methyl 5-ethyl-1-phenyl-1H-1,2,4-triazole-3-carboxylate – Z7957926070 (Method 3).**
Yield: 19%; purity, >95% (assessed by LC/MS).
LC/MS (APSI) m/z [M+H] calculated for $C_{18}H_{22}N_3O_2$: 312.2; found: 312.1.

**3-(2H-1,3-benzodioxol-4-yl)-1-(5-chloro-3,3-dimethyl-2,3-dihydro-1H-indol-1-yl)propan-1-one – Z7957857882 (Method 2).**
Yield: 54%; purity, >95% (assessed by LC/MS).
LC/MS (APSI) m/z [M+H] calculated for $C_{20}H_{21}ClNO_3$: 358.1; found: 358.2.

**(4,5-dihydro-1,2-oxazol-3-yl)methyl 1-(3-fluorophenyl)-4,5,6,7-tetrahydro-1H-indazole-3-carboxylate – Z3646785038 (Method 3).**
Yield: 38%; purity, >95% (assessed by LC/MS).
LC/MS (APSI) m/z [M+H] calculated for $C_{18}H_{19}FN_3O_3$: 344.1; found: 344.2.

**(4R)-3,4-dihydro-2H-1-benzothiopyran-4-yl 4-ethyl-2-(1H-pyrrol-1-yl)-1,3-thiazole-5-carboxylate – Z7957859047 (Method 3).**
Yield: 22%; purity, >95% (assessed by LC/MS).
LC/MS (APSI) m/z [M+H] calculated for $C_{19}H_{19}N_2O_2S_2$: 371.1; found: 371.2.

**2-fluoro-2-methylpropyl 1-(4-fluorophenyl)-4,5,6,7-tetrahydro-1H-indazole-3-carboxylate – Z7957858729 (Method 3).**
Yield: 30%; purity, >95% (assessed by LC/MS).
LC/MS (APSI) m/z [M+H] calculated for $C_{18}H_{21}F_2N_2O_2$: 335.2; found: 335.2.

**(2-fluoro-6-methoxyphenyl)methyl 1-cyclohexyl-5-methyl-1H-pyrazole-3-carboxylate – Z7957859063 (Method 3).**
Yield: 35%; purity, >95% (assessed by LC/MS).
LC/MS (APSI) m/z [M+H] calculated for $C_{19}H_{24}FN_2O_3$: 347.2; found: 347.1.

**[(1R,5R)-bicyclo[3.1.0]hexan-1-yl]methyl 3-phenyl-5H,6H,8H-imidazo[4,3-c][1,4]oxazine-1-carboxylate – Z3171913356 (Method 3).**
Yield: 34%; purity, >95% (assessed by LC/MS).
LC/MS (APSI) m/z [M+H] calculated for $C_{20}H_{23}N_2O_3$: 339.2; found: 339.2.

**[2-methyl-4-(trifluoromethyl)furan-3-yl]methyl 2-methanesulfinylbenzoate – Z3378971266 (Method 3).**
Yield: 50%; purity, >95% (assessed by LC/MS).
LC/MS (APSI) m/z [M+H] calculated for $C_{15}H_{14}F_3O_4S$: 347.1; found: 347.0.

**3-(3-fluoro-2-methoxybenzoyl)-3-azatricyclo[7.3.1.05,13]trideca-1(13),9,11-triene – Z6969215892 (Method 2).**
Yield: 55%; purity, 95% (assessed by LC/MS).
LC/MS (APSI) m/z [M+H] calculated for $C_{20}H_{21}FNO_2$: 326.2; found: 326.1.

**methyl 2-(3-chlorophenyl)-2-(1,2,3,4-tetrahydronaphthalene-1-carbonyloxy)acetate – Z1200201518 (Method 3).**
Yield: 20%; purity, >95% (assessed by LC/MS).
LC/MS (APSI) m/z [M+H] calculated for $C_{20}H_{20}ClO_4$: 359.1; found: 359.0.

**1-{[5-(1-benzothiophen-4-yl)-1,2,4-oxadiazol-3-yl]methyl}-1H-1,3-benzodiazole – Z6969215941 (Method 1).**
Yield: 38%; purity, >95% (assessed by LC/MS).
LC/MS (APSI) m/z [M+H] calculated for $C_{18}H_{13}N_4OS$: 333.1; found: 333.0.

**[(2R)-1-[(4-chloro-3-methyl-1-benzothiophen-2-yl)sulfonyl]-2,3-dihydro-1H-indol-2-yl]methanol – Z6969216135 (Method 7).**
Yield: 45%; purity, >95% (assessed by LC/MS).
LC/MS (APSI) m/z [M+H] calculated for $C_{18}H_{17}ClNO_3S_2$: 394.0; found: 394.0.

**5-(3-chloro-1,2-thiazol-5-yl)-1-(1-methyl-1H-pyrazol-3-yl)-3-[(2-methylphenyl)methyl]-1H-1,2,4-triazole – Z6969216058 (Method 5).**
Yield: 10%; purity, >95% (assessed by LC/MS).
LC/MS (APSI) m/z [M+H] calculated for $C_{17}H_{16}ClN_6S$: 371.1; found: 371.1.

**1-[1-cyclopropyl-3-(pyridin-2-yl)-1H-1,2,4-triazol-5-yl]-5-methylisoquinoline – Z6969216155 (Method 5).**
Yield: 24%; purity, >95% (assessed by LC/MS).
LC/MS (APSI) m/z [M+H] calculated for $C_{20}H_{18}N_5$: 328.2; found: 328.2.

**4-cyclopropyl-2-[1-(2-methylprop-2-en-1-yl)-3-[(pyridin-3-yl)methyl]-1H-1,2,4-triazol-5-yl]pyridine – Z6969216065 (Method 5).**
Yield: 13%; purity, >95% (assessed by LC/MS).
LC/MS (APSI) m/z [M+H] calculated for $C_{20}H_{22}N_5$: 332.2; found: 332.2.

**3-[1-(3-chlorophenyl)-3-[(thiophen-2-yl)methyl]-1H-1,2,4-triazol-5-yl]-4-methyl-1,2,5-oxadiazole – Z6969215903 (Method 5).**
Yield: 12%; purity, >95% (assessed by LC/MS).
LC/MS (APSI) m/z [M+H] calculated for $C_{16}H_{13}ClN_5OS$: 358.1; found: 357.9.

**3-[2-(8-methyl-2,3,4,5-tetrahydro-1,5-benzothiazepin-5-yl)-2-oxoethyl]-1,3-dihydro-2-benzofuran-1-one – Z1745610848 (Method 2).**
Yield: 36%; purity, >95% (assessed by LC/MS).
LC/MS (APSI) m/z [M+H] calculated for $C_{20}H_{20}NO_3S$: 354.1; found: 354.0.

**3-[2-(7-chloro-2,3,4,5-tetrahydro-1,5-benzothiazepin-5-yl)-2-oxoethyl]-1,3-dihydro-2-benzofuran-1-one – Z1355328651 (Method 2).**
Yield: 52%; purity, >95% (assessed by LC/MS).
LC/MS (APSI) m/z [M+H] calculated for $C_{19}H_{17}ClNO_3S$: 374.1; found: 374.2.

**3-[2-(7-chloro-2-methyl-2,3,4,5-tetrahydro-1,5-benzothiazepin-5-yl)-2-oxoethyl]-1,3-dihydro-2-benzofuran-1-one – Z6755334501 (Method 2).**
Yield: 38%; purity, >95% (assessed by LC/MS).
LC/MS (APSI) m/z [M+H] calculated for $C_{20}H_{19}ClNO_3S$: 388.1; found: 388.0.

**3-[2-(7-chloro-3-methyl-2,3,4,5-tetrahydro-1,5-benzothiazepin-5-yl)-2-oxoethyl]-1,3-dihydro-2-benzofuran-1-one – Z1654283767 (Method 2).**

Yield: 44%; purity, >95% (assessed by LC/MS).
LC/MS (APSI) m/z [M+H] calculated for $C_{20}H_{19}ClNO_3S$: 388.1; found: 388.0.

**3-{2-oxo-2-[7-(trifluoromethyl)-2,3,4,5-tetrahydro-1,5-benzothiazepin-5-yl]ethyl}-1,3-dihydro-2-benzofuran-1-one – Z8028935255 (Method 2).**
Yield: 45%; purity, >95% (assessed by LC/MS).
LC/MS (APSI) m/z [M+H] calculated for $C_{20}H_{17}F_3NO_3S$: 408.1; found: 408.0.

**3-{2-oxo-2-[7-(trifluoromethyl)-2,3,4,5-tetrahydro-1,5-benzoxazepin-5-yl]ethyl}-1,3-dihydro-2-benzofuran-1-one – Z8028935249 (Method 2).**
Yield: 75%; purity, >95% (assessed by LC/MS).
LC/MS (APSI) m/z [M+H] calculated for $C_{20}H_{17}F_3NO_4$: 392.1; found: 392.0.

**3-[2-oxo-2-(7-phenyl-2,3,4,5-tetrahydro-1,5-benzoxazepin-5-yl)ethyl]-1,3-dihydro-2-benzofuran-1-one – Z8930854171 (Method 2).**
Yield: 41%; purity, >95% (assessed by LC/MS).
LC/MS (APSI) m/z [M+H] calculated for $C_{25}H_{22}NO_4$: 400.2; found: 400.2.

**3-{2-[7-(5-fluorothiophen-2-yl)-2,3,4,5-tetrahydro-1,5-benzoxazepin-5-yl]-2-oxoethyl}-1,3-dihydro-2-benzofuran-1-one – Z8930854180 (Method 2).**
Yield: 32%; purity, >95% (assessed by LC/MS).
LC/MS (APSI) m/z [M+H] calculated for $C_{23}H_{19}FNO_4S$: 424.1; found: 424.2.

**6-benzyl-3-(2-methoxyphenyl)-[1,2,4]triazolo[3,4-a]phthalazine – Z8144700717 (Method 8).**
Yield: 66%; purity, >95% (assessed by LC/MS).
LC/MS (APSI) m/z [M+H] calculated for $C_{23}H_{19}N_4O$: 367.2; found: 367.2.

**6-benzyl-3-(2-chloro-6-methoxyphenyl)-[1,2,4]triazolo[3,4-a]phthalazine – Z8770784602 (Method 8).**
Yield: 43%; purity, >95% (assessed by LC/MS).
LC/MS (APSI) m/z [M+H] calculated for $C_{23}H_{18}ClN_4O$: 401.1; found: 401.2.

**6-benzyl-3-(2-fluoro-6-methoxyphenyl)-[1,2,4]triazolo[3,4-a]phthalazine – Z8770784612 (Method 8).**
Yield: 11%; purity, >95% (assessed by LC/MS).
LC/MS (APSI) m/z [M+H] calculated for $C_{23}H_{18}FN_4O$: 385.1; found: 385.4.

**6-benzyl-3-(3-chloro-2-methoxyphenyl)-[1,2,4]triazolo[3,4-a]phthalazine – Z8770784603 (Method 8).**
Yield: 52%; purity, >95% (assessed by LC/MS).
LC/MS (APSI) m/z [M+H] calculated for $C_{23}H_{18}ClN_4O$: 401.1; found: 401.0.

**6-benzyl-3-(4-chloro-2-methoxyphenyl)-[1,2,4]triazolo[3,4-a]phthalazine – Z8770784608 (Method 8).**
Yield: 45%; purity, >95% (assessed by LC/MS).
LC/MS (APSI) m/z [M+H] calculated for $C_{23}H_{18}ClN_4O$: 401.1; found: 401.0.

**methyl (1S,3aS,6aR)-2-[4-ethyl-3-(morpholine-4-sulfonyl)benzoyl]-octahydrocyclopenta[c]pyrrole-1-carboxylate – Z9108382167 (Method 2).**
Yield: 52%; purity, >95% (assessed by LC/MS).
LC/MS (APSI) m/z [M+H] calculated for $C_{22}H_{31}N_2O_6S$: 451.2; found: 451.2.

**(1R)-1-[3-(5-{3-bromopyrazolo[1,5-a]pyridin-4-yl}-1,2,4-oxadiazol-3-yl)phenyl]ethan-1-ol – Z6969215882 (Method 1).**
Yield: 29%; purity, >95% (assessed by LC/MS).
LC/MS (APSI) m/z [M+H] calculated for $C_{17}H_{14}BrN_4O_2$: 387.0; found: 387.0.

**3-(methylsulfanyl)phenyl 3-benzyl-1-methyl-1H-pyrazole-4-carboxylate – Z6969215899 (Method 2).**
Yield: 36%; purity, >95% (assessed by LC/MS).
LC/MS (APSI) m/z [M+H] calculated for $C_{19}H_{19}N_2O_2S$: 339.1; found: 339.0.

**3-[(2,3-dihydro-1-benzofuran-3-yl)methyl]-5-(4-methylfuran-2-yl)-1-(2-methylprop-2-en-1-yl)-1H-1,2,4-triazole – Z6969216160 (Method 5).**
Yield: 28%; purity, >95% (assessed by LC/MS).
LC/MS (APSI) m/z [M+H] calculated for $C_{20}H_{22}N_3O_2$: 336.2; found: 336.2.
